# Ratiometric Quantification of Dissolved Molecular Oxygen in Microplates for Biochemical Assays Using Palladium Porphyrin Photoluminescence

**DOI:** 10.64898/2026.04.15.718663

**Authors:** Dina Podolskiy, Christoph Plieth

## Abstract

Many biochemical processes are dependent on the presence or absence of molecular oxygen (O_2_). Palladium-tetrapyrrol derivatives can be used to measure O_2_-concentrations and O_2_-turnover during biochemical reactions and microbial growth in standard microtiter plates (MTPs). Palladium(II)-5,10,15,20-(tetrapentafluorophenyl)-porphyrin (**1**; CAS 72076-09-6) and Palladium(II)-5,10,15,20-(tetraphenyl)tetrabenzoporphyrin (**2**; CAS 119654-64-7) are introduced with this study. Spectral analyses of both compounds revealed that fluorescence quenching by O_2_ is not evenly distributed throughout all wavelengths and can therefore be used ratiometrically. Experimentally determined fluorescence lifetimes are around 500 µs and 300 µs for **1** and **2**, respectively. A simple protocol is disclosed, how to immobilize the indicators on the bottom of MTP wells to give clear transparent dye doped polymer layers. We propose a straightforward procedure of how fluorescence data can be processed and calibrated in terms of O_2_ concentrations. Diverse applications are demonstrated and discussed, which include oxygen consumption and production by microorganisms as well as by enzymatically catalysed biochemical reactions. Various aspects are critically considered, as there are e.g. the dependence of O_2_ solubility on temperature and salinity, the diffusion of O_2_ across diverse phase boundaries, the unwanted O_2_ ingress into the reaction volume, the oxygen binding capacity of the MTP plastic material and the pH-dependence of the sensor layer. The findings and methods presented here open up a broad variety of high throughput assays involving changes of dissolved O_2_ as measurands for biochemical and biological activity.

**Graphical Abstract:** 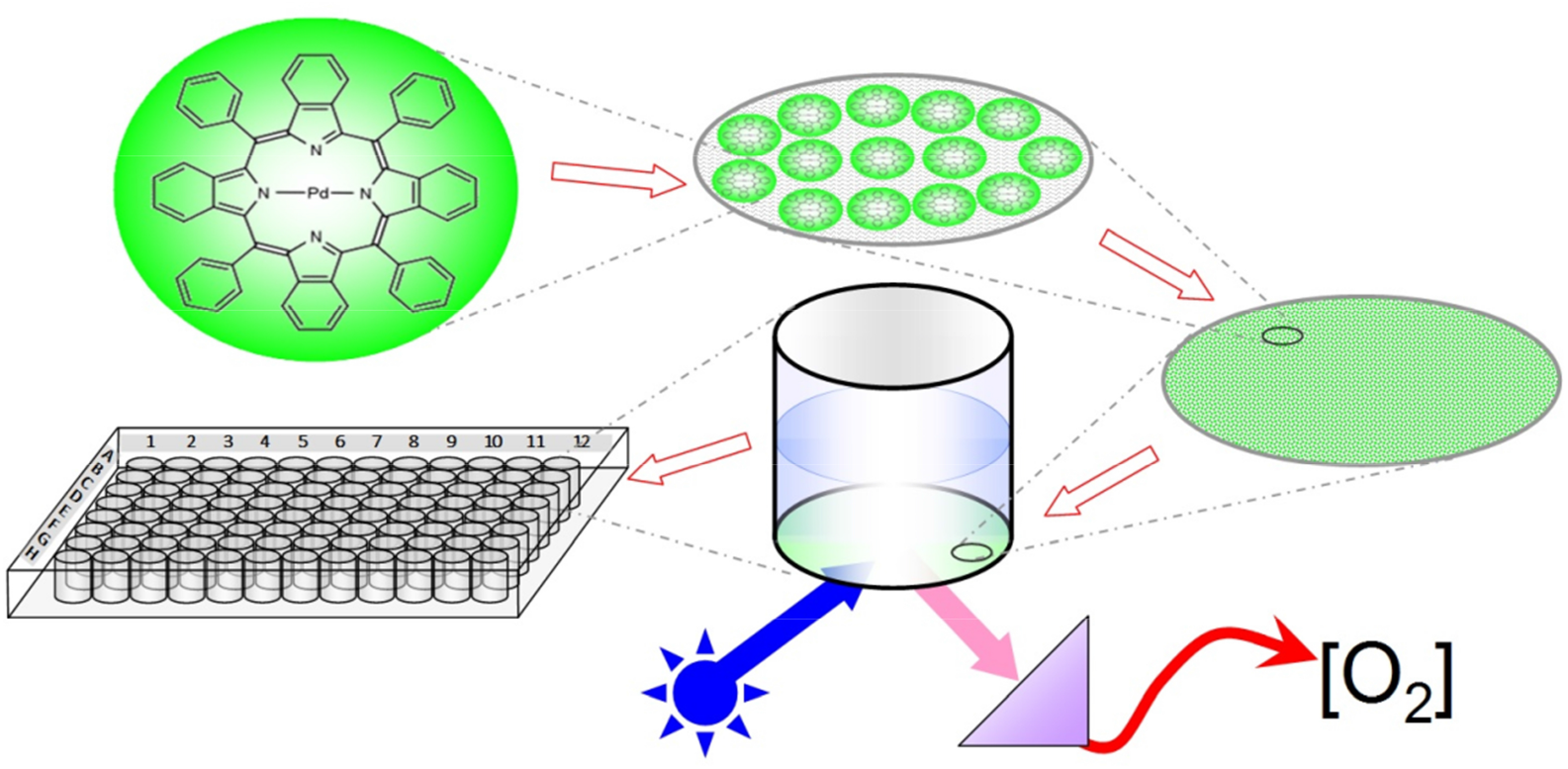

## Introduction

All life on earth is dependent on O_2_, since abundant molecular oxygen (O_2_) influences almost all natural processes on our planet’s surface. Nearly 21% of our planet’s troposphere is O_2_, most of which has been generated by photosynthesis. It dissolves in water to a certain extent but also binds to and penetrates through diverse solid materials and surfaces.

In metabolism, O_2_ serves as an electron acceptor and is central for many biochemical redox processes. Diverse biochemical reactions consume or produce O_2_. A prerequisite for this is that it dissolves in aqueous milieu. However, the solubility of oxygen in water is limited to concentrations of [O_2_] < 1 mM (< 30 mg/ml; details see DOTABLES ^1–3^).

Oxygen consumption and production by living organisms are good indicators for metabolic processes like respiration ^4^ and photosynthesis ^5–7^, respectively. Studying these processes requires precise methods to quantify [O_2_] and changes thereof in real time. In life science laboratories, two popular techniques are mainly in use. Both have advantages and limitations ^8,9^.

First, amperometric electrodes exploit the effect that electrical currents, conducted by electrolyte solutions, are dependent on [O_2_] ^6^. Clark-type oxygen electrodes are widely used in life science laboratories, as well as with water quality monitoring and field measurements ^10–12^. The output signal is linearly correlated with [O_2_] ^13^. Hence, a simple two-point calibration can be used to convert the amperometric signal into [O_2_].

Second, the fluorescence of specific molecules can be quenched by O_2_ ^14^. Devices exploiting this effect for [O_2_] quantification are called optodes or optrodes and are used with many biosensor applications ^9,15–17^. Noble metal porphyrins (NMPs) are widely used for these purposes ^18–22^. However, the development and production of specific biosensor opt(r)odes is not routine and requires some experience and engineering skills ^23–31^.

Nowadays, microtiterplate (MTP) readers recording optical signals like absorbance, fluorescence or chemiluminescence, belong to the standard equipment of many life science laboratories for high throughput (HTP) quantification of diverse parameters. Consequently, specific MTP assays have been developed, which also allow the quantification of [O_2_] on the basis of fluorescence quenching ^32,33^.

In particular, fluorescent noble metal porphyrins (NMPs) are on the market for quantification of [O_2_] with fiber optics or in MTPs ^9,21,34^ . However, due to some frame parameters, these optrode products are not widely established. They are costly and some limitations such as undesirable oxygen ingress require attention and control experiments, which, in turn, require more of the material.

The properties of any O_2_-optrode depends mainly on the quenching effect caused by O_2_, which depends on several factors such as fluorescence lifetime, temperature, diffusion rate of O_2_ within the sensor matrix and within the aqueous sample to be probed. In addition, the calibration, i.e. the conversion of fluorescence signals from O_2_ optrodes into precise values of dissolved oxygen concentrations ([O_2_]), needs some thoughts as O_2_ solubility depends on factors like temperature, salinity, and barometric pressure (**Supporting Information**: Temperature dependence of O_2_ solubility **Fig SI 1**, Salinity dependence of O_2_ - solubility **Fig SI 2**) ^1^ .

The two indicators introduced here (**Fig. 1**) are in the following named Pd-Fluorophorphyrin (**1**) and Pd-Benzophorphyrin (**2**) for short. We present the relevant spectral characteristics of **1** and **2**, and provide a simple protocol how to immobilize them in standard transparent polystyrene microtiter plates (PS-MTPs) for ratiometric quantification of [O_2_], using the fluorescence bottom-read option. We propose a straightforward calibration procedure to calculate [O_2_] from ratiometric fluorescence data, thereby taking the dependence of O_2_ solubility on temperature and salinity into account.

**Figure 1.**
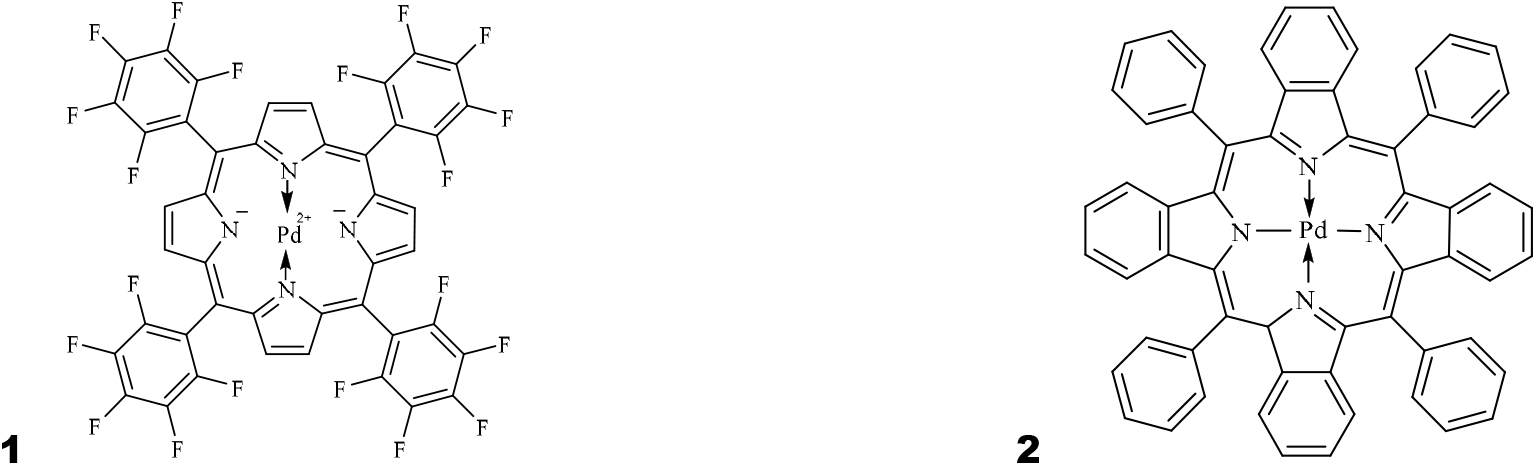
Structures of noble metal porphyrins (NMPs) used as O_2_ indicators for experiments presented below **1** Palladium(II)-5,10,15,20-(tetrapentafluorophenyl)porphyrin (CAS 72076-09-6) MW = 1079 g/mol **2** Palladium(II)-5,10,15,20-(tetraphenyl)tetrabenzoporphyrin (CAS 119654-64-7) MW = 919 g/mol

As a proof of concept we demonstrate, how specific biochemical reactions involving consumption or production of O_2_ can be characterized. In addition, we employed bacteria and microalgae to show how their consumption and production of O_2_ can be quantified under diverse conditions. Finally, we provide in detail information on how to deal with problematic frame parameters, such as e.g. undesired O_2_ ingress.

## Results & Discussion

### Absorbance spectra and daylight appearance of the O_2_-indicators

Absorbance spectra (Fig. 2) of both NMPs (**1** and **2**; Fig. 1) appear as typical metalloporphyrin spectra with Soret-band in the blue region and two Q-Bands at longer wavelengths, namely the vibronic (Q_1_) and the non-vibronic band (Q_0_) ^35–37^. Key parameters of both spectra are summarized in **Tab. 1**.

**Table 1.** Absorbance Characteristics.

|  | Pd-Fluoroporphyrin<br>**(1)** | Pd-Benzoporphyrin<br>**(2)** |
| --- | --- | --- |
| Soret-band peak (FWHM) | 412 nm (26 nm) | 446 nm (20 nm) |
| Vibronic Q<sub>1</sub>-band peak (FWHM)<br>Peak height relative to Soret | 520 nm (27 nm)<br>0.10 | 582 nm (30 nm)<br>0.07 |
| Non-vibronic Q<sub>0</sub>-band peak (FWHM)<br>Peak height relative to Soret | 554 nm (16 nm)<br>0.08 | 632 nm (22 nm)<br>0.47 |

**Figure 2.**
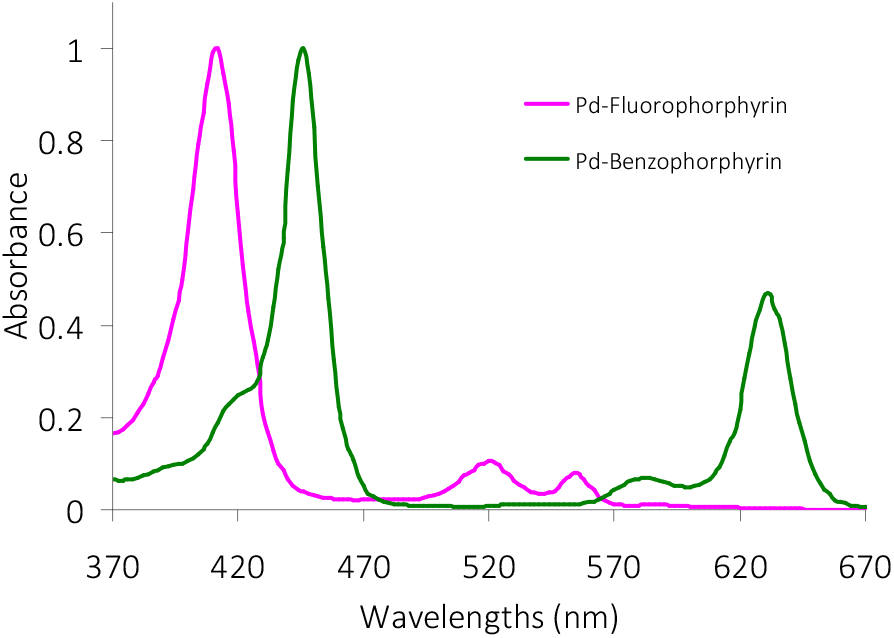
Absorbance spectra of 1 and 2. MTP wells were doped with Pd-Porphyrins as described in the experimental section. Absorbance of the indicator layer was scanned by a plate reader and spectra were normalized by the peak of their Soret band. Pd-Fluoroporphyrin (**1**) absorbs mainly in the blue and green regions an thus appears red, while Pd-Benzoporphyrin (**2**) absorbs in the blue and the red region, having a distinct green absorption gap and thus appears, like chlorophyll, green. Spectral characteristics of the peaks are summarized in **Tab. 1**

Both, **1** and **2**, absorb visible light. **1** appears red-pink (**Supporting Information: Fig. SI 1** Flat bottom MTPs for [O_2_] quantification) because it absorbs mainly in the blue region around 412 nm (**Fig. 2**), whereas **2** has two absorbance peaks in the visible range at 446 nm and 632 nm and thus, like chlorophyll, looks green (**Fig. SI 1**). The main difference in colour is caused by a lack of absorbance in the 600 nm-range as seen in the spectrum of **1** (**Fig. 2**).

### Photoluminescence Spectra

The excitation spectra of **1** and **2** (**Figs. 3A, 4A**) are similar in shape when compared with their absorption spectra (**Fig. 2**), indicating that absorbed light is converted to fluorescence at all wavelength and heat dissipation (i.e. vibronic excitation) is negligible. However, the indicators’ O_2_ sensitivity is not evenly distributed throughout all wavelengths. This is demonstrated by the ratio spectra (black lines in **Figs. 3** and **4**). The ratio of fluorescence emission intensities from O_2_-low and O_2_-rich wells varies and reaches values of > 4. Higher ratios are not reached here, due to the incomplete deprivation of [O_2_] by Na_2_SO_3_ and the temperature dependent limited O_2_ solubility in aqueous solutions (**Supporting information: Fig. SI 2** Temperature dependence of O_2_ solubility).

**Figure 3.**
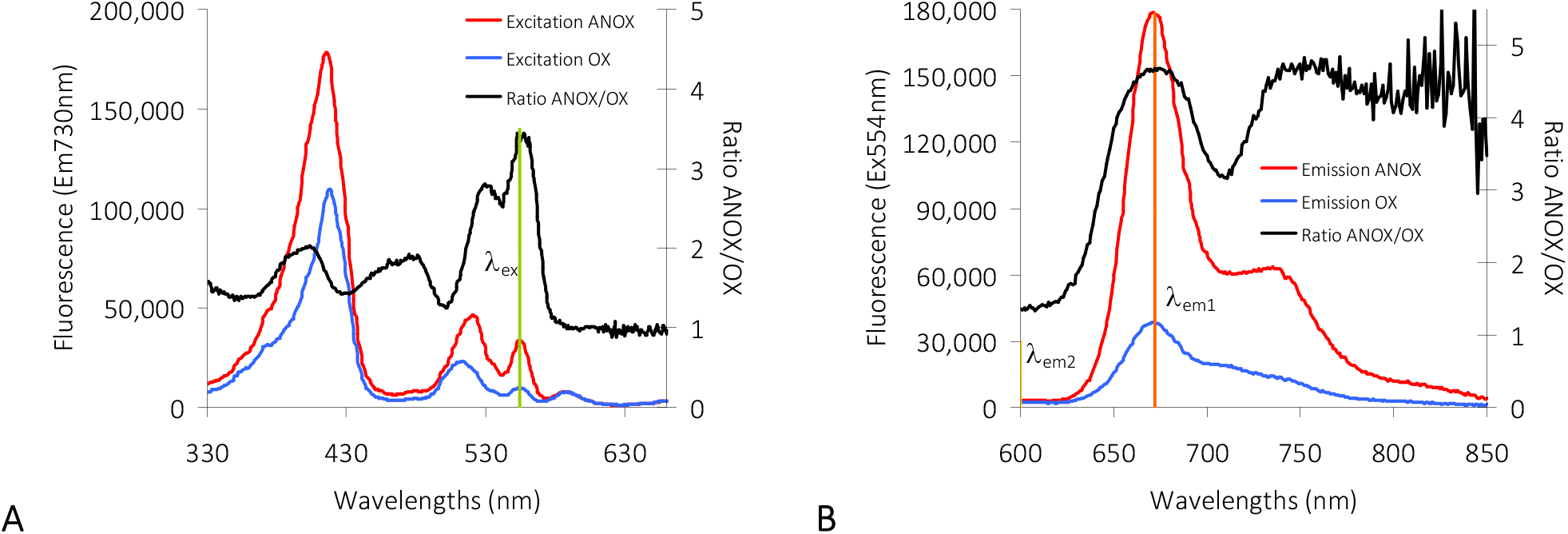
Fluorescence spectra of Pd-Fluorophorphyrin. (**1**) under oxygenated conditions (blue lines; wells filled with H_2_O) and anoxic conditions (red lines; wells filled with 2 M Na_2_SO_3_ solution). Black lines indicate the ratio of the ANOX-data divided by the OX-data calculated for each wavelength. Fluorescence is given in arbitrary units. Spectra are averages of n = 12 replicates. **A**: Excitation spectra taken with emission at λ_em_ = 730nm; **B**: Emission spectra taken with excitation at λ_ex_ = 554 nm.

**Figure 4.**
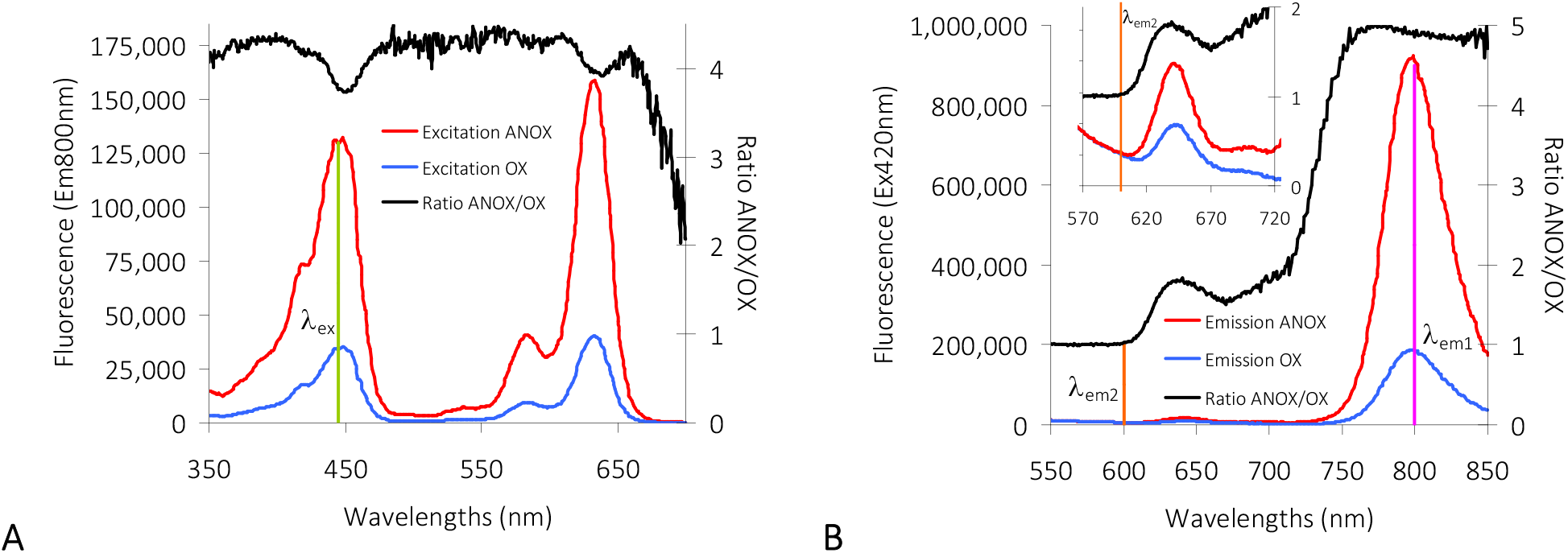
Fluorescence spectra of Pd-Benzophorphyrin (2) under oxygenated conditions (blue lines; wells filled with H_2_O) and anoxic conditions (red lines; wells filled with 2 M Na_2_SO_3_ solution). Black lines indicate the ratio of the ANOX-data divided by the OX-data calculated for each wavelengths. Fluorescence is given in arbitrary units. Spectra are averages of n = 12 replicates. **A**: Excitation spectra taken with emission at λ_em_ = 800 nm; **B**: Emission spectra taken with excitation at λ_ex_ = 420 nm.

### Characteristics of fluorescence spectra

The recognition of wavelength dependent quenching and of spectral shifts upon quencher collision or ligand binding is possible when spectra are normalized by their area. If the quenching effect was evenly distributed, then normalized spectra taken under O_2_ saturated conditions (OX) would be identical with normalized spectra taken under low [O_2_] (ANOX). Hence, the difference between both normalized spectra (designated “MiniMax spectrum”; **Fig. 5**) would be constant null. However, as shown in **Fig. 5**, the normalized emission spectra reveal wavelength dependent quenching.

When fluorescent indicators are quenched equally at all wavelengths, then a quencher-independent fluorescer is required as internal standard for (pseudo-)ratiometric measurements. For Pt-Porphyrin as oxygen indicator, Rhodamine was used as a reference standard to enable ratiometry ^21,38,39^. However, when fluorescence quenching is not equally distributed over all wavelengths, as is the case here (**Fig. 5A** and **B**), then two different fluorescent species could be assumed in the sample. The spectrum obtained is then an overlay of their distinct spectra. This effect becomes evident, with the data set shown in **Fig. 5**. The discontinuous minima (null points) of the difference spectrum (‘MiniMax’; grey lines **Fig. 5**) reveal the isoemission points. These iso-points separate the wavelength range for measuring the desired quantity, here [O_2_], from wavelengths that can serve as an internal reference. They make evident that emission intensities at different wavelengths can be used for ratiometric measurements.

### Selecting appropriate wavelengths for [O_2_] measurements

Finding the optimal wavelengths for [O_2_] measurements requires a compromise over maximum fluorescence intensity and oxygen sensitivity.

With Pd-Fluoroporphyrin (**1**) the maximum O_2_ sensitivity in excitation is given with the Q_0_-band at 554 nm (**Fig. 3 A**), although maximum fluorescence can be achieved with excitation in the Soret band around λ_ex_ = 412 nm. Considering fluorescence emission, peak fluorescence and maximum O_2_ sensitivity are at λ_em1_ = 672 nm (**Fig. 3 B**). At λ_em2_ = 600 nm emission fluorescence is almost O_2_ insensitive (Ratio ANOX/OX ≈ 1). This suggests using fluorescence at this emission wavelength as reference for ratioing (**Tab. 2**).

**Table 2.** Excitation and emission wavelengths. The fluorescence intensity ratios R = F_λem1_/F_λem2_ were used as indicator variable for dissolved O_2_. Wavelengths have been chosen according to fluorescence yields and O_2_ responsiveness and are used for all experiments described in this study. The numerator wavelength λ_em1_ for ratioing is at maximum emission. The denominator wavelength λ_em2_, is where low or no fluorescence quenching is seen (i.e. Ratio ANOX/OX ≈ 1; **Figs. 3 B, 4 B**)

| The fluorescence intensity ratios $R = F_{\lambda_{em1}}/F_{\lambda_{em2}}$ were used as indicator variable for dissolved O<sub>2</sub>. Wavelengths have been chosen according to fluorescence yields and O<sub>2</sub> responsiveness and are used for all experiments described in this study. The numerator wavelength $\lambda_{em1}$ for ratioing is at maximum emission. The denominator wavelength $\lambda_{em2}$ is where low or no fluorescence quenching is seen (i.e. Ratio ANOX/OX $\approx 1$; **Figs. 3 B, 4 B**) | | | | | |
| --- | --- | --- | --- | --- | --- |
| Pd-Fluoroporphyrin (**1**) |  |  | Pd-Benzophorphyrin (**2**) |  |  |
| $\lambda_{ex}$ | $\lambda_{em1}$ | $\lambda_{em2}$ | $\lambda_{ex}$ | $\lambda_{em1}$ | $\lambda_{em2}$ |
| 554 nm | 672 nm | 600 nm | 445 nm | 800 nm | 600 nm |

The situation is different with Pd-Benzophorphyrin (**2**). Maximum fluorescence emission and maximum O_2_ sensitivity are around λ_em1_ = 800 nm (**Fig. 4 B**). O_2_ insensitivity with still reasonable fluorescence intensity is seen with λ_em2_ = 600. When choosing these two wavelengths for ratioing, neither the Q_0_ nor Q_1_ band can be used for excitation. Hence excitation in the Soret-band at λ_ex_ = 445nm is required to achieve acceptable fluorescence intensities (**Fig. 4 A**). The wavelengths for fluorescence emission ratioing (R = F_λ em1_/F_λem2_) and [O_2_] calculation chosen for all further experiments described below are summarized in **Tab. 2**.

**Figure 5.**
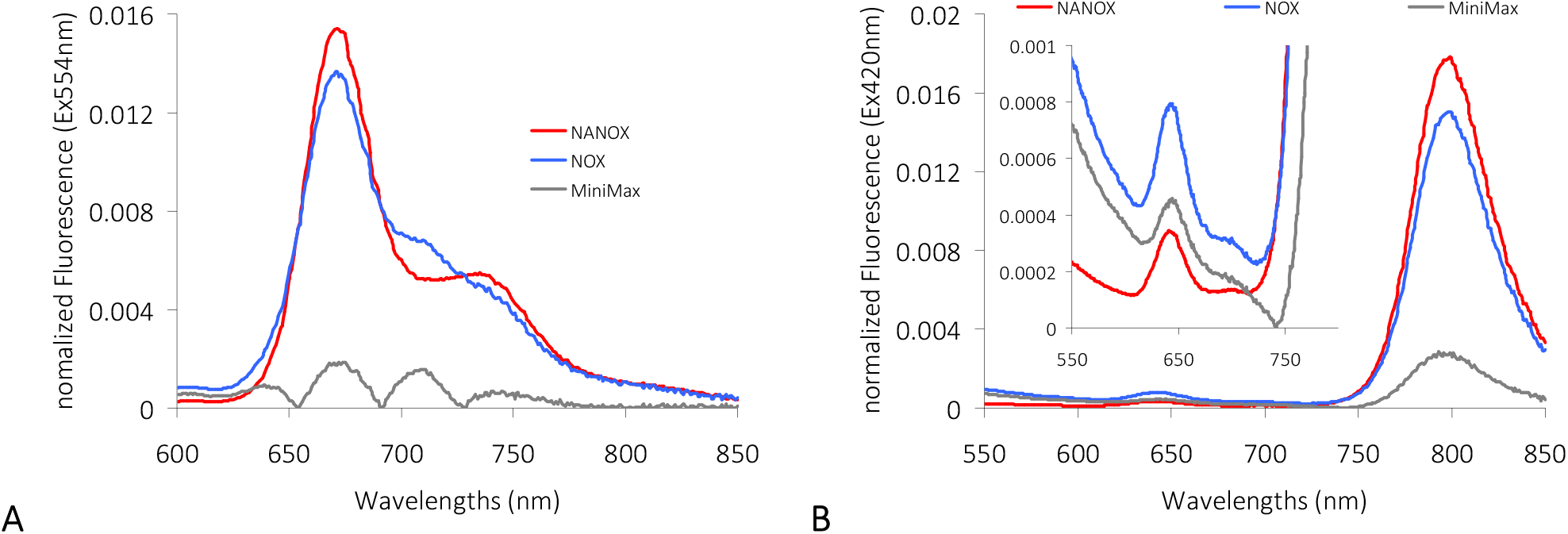
Fluorescence emission spectra normalized by the area under the curve (A = 1). MiniMax spectra (grey lines) are the absolute difference between ANOX and OX **A:** normalized spectra of **1** calculated from spectra shown in **Fig. 3B**. The nulls of the MiniMax spectrum reveal isoemission (‘isosbestic’) points at 655 nm, 692 nm and 729 nm **B**: normalized spectra of **2** calculated from spectra in **Fig. 4B**. An isoemission point obtained from null of MiniMax is at 744 nm. All spectra are averages of n = 12 technical replicates.

### Fluorescence lifetimes and adjustment of integration times

The choice of an appropriate integration time T for photoluminescence measurements is important, because it affects both, the maximum possible sampling rate and the signal-to-noise ratio. After excitation, the population of excited states decays exponentially and gives off light with a lifetime (i.e. time constant) of τ according to **Eq. 1**.

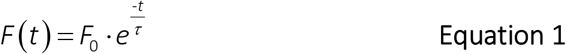

Fluorescence lifetimes τ of NMPs are typically in the range between 20 µs and 2000 µs ^18,40–43^. Direct lifetime measurements are not possible with a plate reader. However, the instrument allows to precisely set the duration of photon counting T (i.e. integration time) and thereby perform a multi-windows method ^43^. Thus, NMP fluorescence photons collected with various integration times T at fixed gain settings provides means of measuring the lifetime τ. The amount of light emitted during the decay process (**Eq. 1**) is determined by T (**Eqs. 2**).

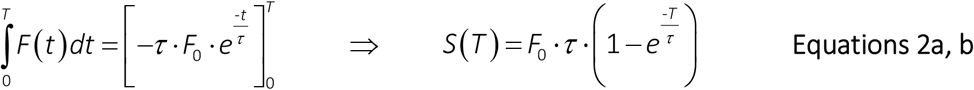

This integral S(T) is a function of T with lifetime τ as a parameter (**Eq. 2b**).

Series of integration times T from 20 to 2000 µs gave data sets S(T) as shown in **Fig. 6**. Fitting Chapman model functions (**Eq. 3**) to such data with the exponent c fixed to c = 1 revealed parameters a and b.

**Figure 6.**
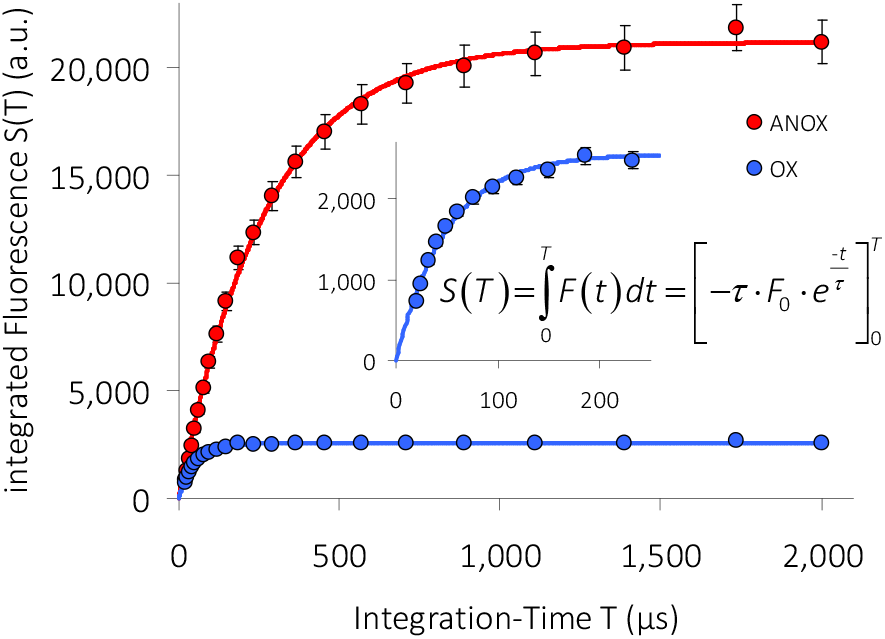
Integrated fluorescence S from **2** plotted over integration time T demonstrates how fluorescence life times τ under anoxia *versus* oxygenated conditions were obtained. A series of fluorescence S (λ_ex_ = 445nm; λ_em_ = 800nm) was collected with increasing integration times T (20 µs to 2000 µs). MTP wells were filled either with O_2_ absorber and sealed (red line; oxygen-free samples) or with H_2_O and left open (blue line; oxygenated samples). A Chapman model function was fitted to the data to obtain the parameters giving S(T). Life times τ were calculated as described in the main text (**Eq. 2-4**). Data are averages of n = 12. Error bars represent SD. SD is below symbol size where error bars cannot be seen.

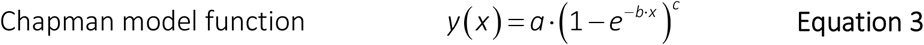

From parameters a and b the parameters τ and F_0_ of S(T) could be calculated (**Eqs. 4a, b**)

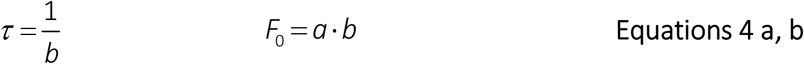

This way, lifetimes τ of **1** and **2** were determined as listed in **Tab. 3**.

**Table 3.** Fluorescence lifetimes τ. **Lifetimes** τ are calculated from integrated fluorescence measurements S(T) (**Eq. 2b**). Values given are averages of n ≥ 5 experiments. Numbers (±) in parentheses denote SD.

| Lifetimes $\tau$ are calculated from integrated fluorescence measurements $S(T)$ (Eq. 2b). Values given are averages of $n \geq 5$ experiments. Numbers (±) in parentheses denote SD. | | | | | | |
| --- | --- | --- | --- | --- | --- | --- |
|  | Pd-Fluorophorphyrin (**1**) |  |  | Pd-Benzophorphyrin (**2**) |  |  |
| Temp. (°C) | $\tau$ ANOX (O<sub>2</sub>-free) | $\tau$ OX (O<sub>2</sub>-saturated) | Stern-Volmer Quotient $\tau_0/\tau$ | $\tau$ ANOX (O<sub>2</sub>-free) | $\tau$ OX (O<sub>2</sub>-saturated) | Stern-Volmer Quotient $\tau_0/\tau$ |
| 25 | 503 $\mu\text{s}$ ($\pm 28$) | 54 $\mu\text{s}$ ($\pm 2.3$) | 9.35 ($\pm 0.5$) | 312 $\mu\text{s}$ ($\pm 13$) | 48 $\mu\text{s}$ ($\pm 1.4$) | 6.55 ($\pm 0.15$) |
| 30 | 510 $\mu\text{s}$ ($\pm 39$) | 52 $\mu\text{s}$ ($\pm 3$) | 9.88 ($\pm 0.78$) | 294 $\mu\text{s}$ ($\pm 16$) | 47 $\mu\text{s}$ ($\pm 1$) | 6.31 ($\pm 0.24$) |
| 37 | 471 $\mu\text{s}$ ($\pm 66$) | 48 $\mu\text{s}$ ($\pm 3$) | 9.89 ($\pm 1.01$) | 295 $\mu\text{s}$ ($\pm 4$) | 45 $\mu\text{s}$ ($\pm 2$) | 6.55 ($\pm 0.2$) |

The data (**Tab. 3**) show that lifetimes are shorter in presence of oxygen, proving dynamic quenching. Values are in the same range as reported elsewhere ^18,40–43^. Thus, for NMPs, integration times in the four-digit range of µs should be chosen for fluorescence recordings. For all further experiments presented here, the platereader was set to an integration time T of 1000 µs.

The rate of collisional quenching typically increases with increasing temperatures. This effect should become visible through a decrease of τ. Here, experiments were performed at different temperatures (25°C; 30°C and 37°C) but gave no significant differences in τ (**Tab. 3**). This is understandable, because the fluorescent indicators are immobilised in a PS layer and the diffusion of oxygen into and within the plastic material is limited when compared with its mobility in aqueous solutions or in air ^44–46^.

### Quantification of oxygen ingress

Aqueous solutions in contact with ambient atmosphere typically have concentrations of few hundred micromolar of dissolved oxygen (100 µM <[O_2_] < 500 µM) when in equilibrium (**Supporting information: Fig. SI 2**; DOTABLES) ^3^. O_2_ can diffuse from the gas phase into the solution (ingress) and *vice versa* (egress). In presence of sulfite (Na_2_SO_3_), dissolved oxygen is consumed and sulfite is converted to sulfate (Na_2_SO_4_) (**Eq. 5**). Cobalt, as a catalyst, accelerates this process ^47^. Thus, a solution with a sufficiently high Na_2_SO_3_ concentration and a few ppm of CoCl_2_ is low in [O_2_].

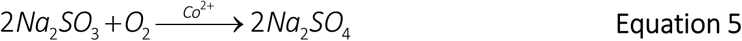

Wells with immobilized Pd-Porphyrins were filled with sulfite solutions of diverse concentrations ([Na_2_SO_3_]). Pd-Porphyrin fluorescence was recorded over several hours during which the wells remained unsealed, allowing O_2_ ingress from the atmosphere.

Quenching of fluorescence F by O_2_ starts as soon as all sulfite is converted to sulfate (**Fig. 7 A**). The derivations of the data (dF/dt) indicate at which time the entire amount of Na_2_SO_3_ in the wells has been consumed by ingressed O_2_ (**Fig. 7 B**).

**Figure 7:**
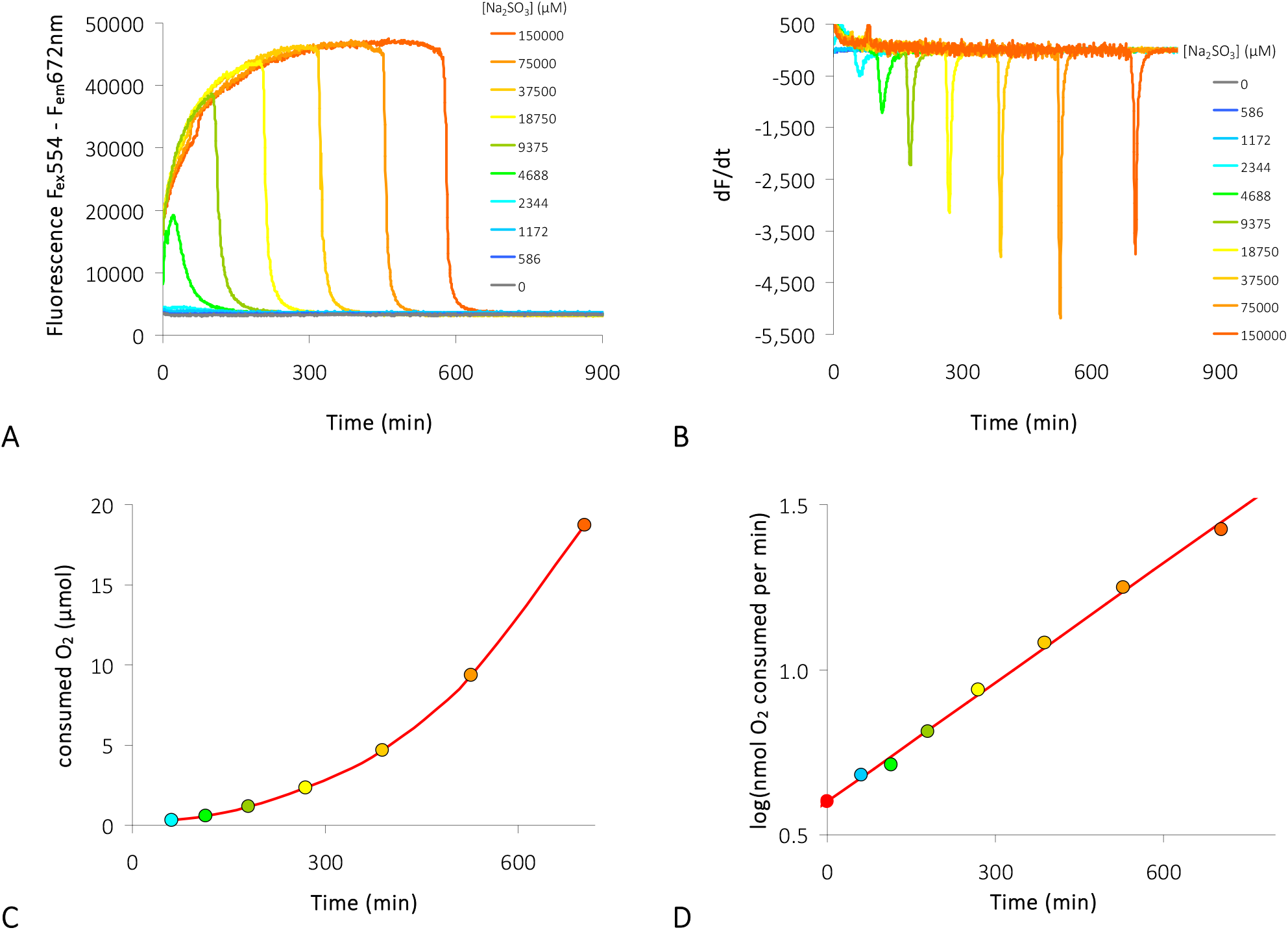
Oxygen ingress quantified using dilution series of Na_2_SO_3_. **A:** Pd-Fluoroporphyrin (**1**) coated MTP loaded with 200 µl of diverse concentrations of Na_2_SO_3_ was exposed to ambient air and fluorescence (λ_ex_ = 554; λ_em_ = 672 nm) was recorded. Sample rate = 1·min^-1^ **B:** The time derivatives calculated from data presented in **A** reveal with their downward peaks the time points (i.e. consumption times), where all Na_2_SO_3_ (sulfite) is converted to Na_2_SO_4_ (sulfate). **C:** The consumed amount O_2_ (in µmol) as calculated from converted amount of Na_2_SO_3_ (**Eq. 5**) plotted over time reveals an exponential dependence. **D:** log of oxygen consumption (in nanomol O_2_ per time (min) calculated from data shown in **C**) plotted over time gives a straight line. Extrapolation of the line intersects with the ordinate (red dot at y = 0.6). This intersection marks the oxygen ingress/egress equilibrium (here 4 nmol/min) if no Na_2_SO_4_ was in the solution.

The amount of O_2_ needed to convert the sulfite can be calculated from the amount of sulfite, the well was loaded with (stoichiometry: 1 O_2_ per 2 Na_2_SO_3_; **Eq. 5**). The consumed amount of O_2_, plotted against time, reveals an exponential increase (**Fig. 7 C**). Consequently, in this process the consumed O_2_ and time are not linearly correlated but rather follow a logarithmic dependence. Thus, in a half-logarithmic plot of log(consumed O_2_/time) over time, a straight line is obtained (**Fig. 7 D**). The extrapolation of the trend intersects with the ordinate. The intersection point reveals the oxygen ingress if there was no Na_2_SO_3_ in solution. This gives the O_2_ ingress rate, which, at equilibrium, is equal to O_2_ egress.

At first glance, a doubling of [Na_2_SO_3_] should lead to a doubling of the time needed to convert all sulfite into sulfate. However, the whole process is driven by diffusion which, in turn, is dependent on concentration gradients (Fick’s laws). The high diffusion rate of O_2_ in the gas phase leads to a quick depletion of sulfite at the solution surface. With increasing [Na_2_SO_3_], steeper vertical concentration gradients of Na_2_SO_3_ are established in the solution. This causes accelerated diffusion in the liquid phase, disproportionately faster turnover (Eq. 5), and shorter consumption times t_c_.

### Proof of the calibration linearity

Since dynamic fluorescence quenching is assumed as described by a Stern-Volmer equation ^48^, a uniform quenching should be expected at all fluorescence wavelengths. However, the NMPs (**1**, **2**) used here show a wavelength-dependent quenching by O_2_ (**Figs. 3, 4**). This raises the speculation that there is no pure dynamic quenching process at work, which may possibly result in a non-linear (e.g. logistic or sigmoidal) relationship between fluorescence ratios R and [O_2_].

We addressed this concern and recorded [O_2_] amperometrically in parallel to the ratiometric method (**Supporting information: Fig. SI 4** Correlating NMP fluorescence ratios with amperometric oxygen recording). An amperometric signal is always linearly correlated with [O_2_], provided the O_2_-electrode is properly prepared ^10,49^. If a non-linear relationship was given between NMP fluorescence ratio R and [O_2_], then this should become visible when comparing the optical signal R with the simultaneously recorded amperometric signal. Since the data confirm linearity (**Supporting information: Figs. SI 4 B, C** - insets), it becomes obvious that the used approach of linear calibration used here (**Eqs. 8 – 11** in the Methods section) is appropriate.

### Adjustments for temperature

The temperature dependence of O_2_ solubility needs to be considered, when [O_2_] is recorded with varying temperature. During fluorescence recording, the plate reader logs the temperature (in °C) inside the instrument. These temperature data can be taken for calibration correction according to Eq. 10 (Methods section below).

When calibration is performed without temperature correction (**Eq. 8**), then a constant [O_2_] at all temperatures probed is the result (**Fig. 8** light symbols). However, this does not correspond to the actual [O_2_] which decreases with increasing temperatures due to the decreased O_2_ solubility^3^ (expressed as [O_2_]_max_) and the resulting O_2_ egress. Thus, with variations in temperature during the experiment, a temperature-dependent correction as given by **Eq. 10** is necessary and leads to more reliable results (**Fig. 8** dark symbols).

**Figure 8.**
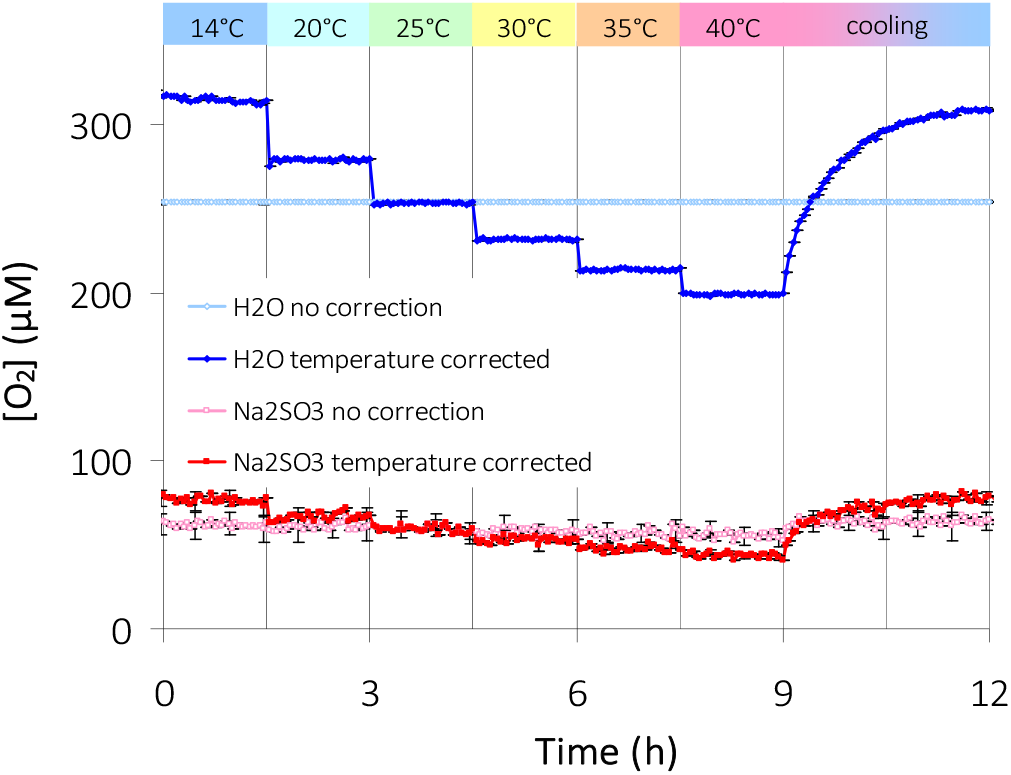
Necessity of temperature correction with ratiometric [O_2_] recording. Wells were filled either with plain water (blue symbols) to achieve [O_2_]_max_ or with a sulfite solution to adjust to a relatively low [O_2_] (red symbols). [O_2_] appears to be constant at all temperatures (light symbols) when fluorescence ratios were converted to [O_2_] using **Eq. 8**, with [O_2_]_max_ = 254 µM (at 25°C), thereby disregarding the temperature dependence of O_2_ solubility. Implementing the temperature correction of [O_2_]_max_ (**Eq. 10**) leads to reliable results (dark symbols) showing that [O_2_] decreases with increasing temperature, because reduced solubility causes O_2_ egress. Experimental conditions: 200 µl per well; sample rate = 1/3 ·min^-1^. [Na_2_ SO_3_ ] = 2M; R_max_ was taken from wells filled with O_2_-absorber powder. Average of n = 12; Error bars represent SD.

### Quantification of O_2_ consumption and production

Some enzymes such as oxygenases or laccases consume O_2_, when catalysing a reaction ^50–53^. Their enzymatic activities can be quantified by monitoring the [O_2_] in the reaction volume ^15,54^. Consequently, NMP fluorescence in MTPs should be useful to characterise such enzymes, to detect, and to quantify their substrates, or to investigate O_2_ consumption or production of whole microorganisms.

### O_2_ consumption by the laccase reaction

The laccase reaction presented here is an example for enzymatically catalysed O_2_ consumption. The enzyme converts pyrogallol to pupurogallin (**Eq. 6**) with an optimum at pH = 5.5 ^55^.

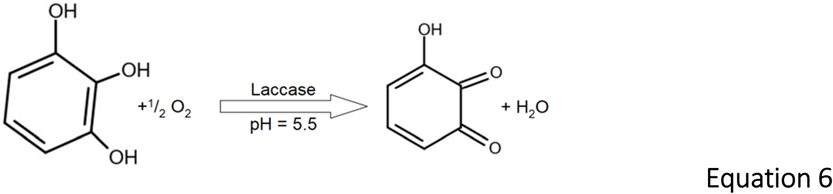

The data presented in **Fig. 9** reveal that a characterization of enzyme turnover is possible. However, interfering processes, such as O_2_ ingress, need to be kept in check (details below).

**Figure 9.**
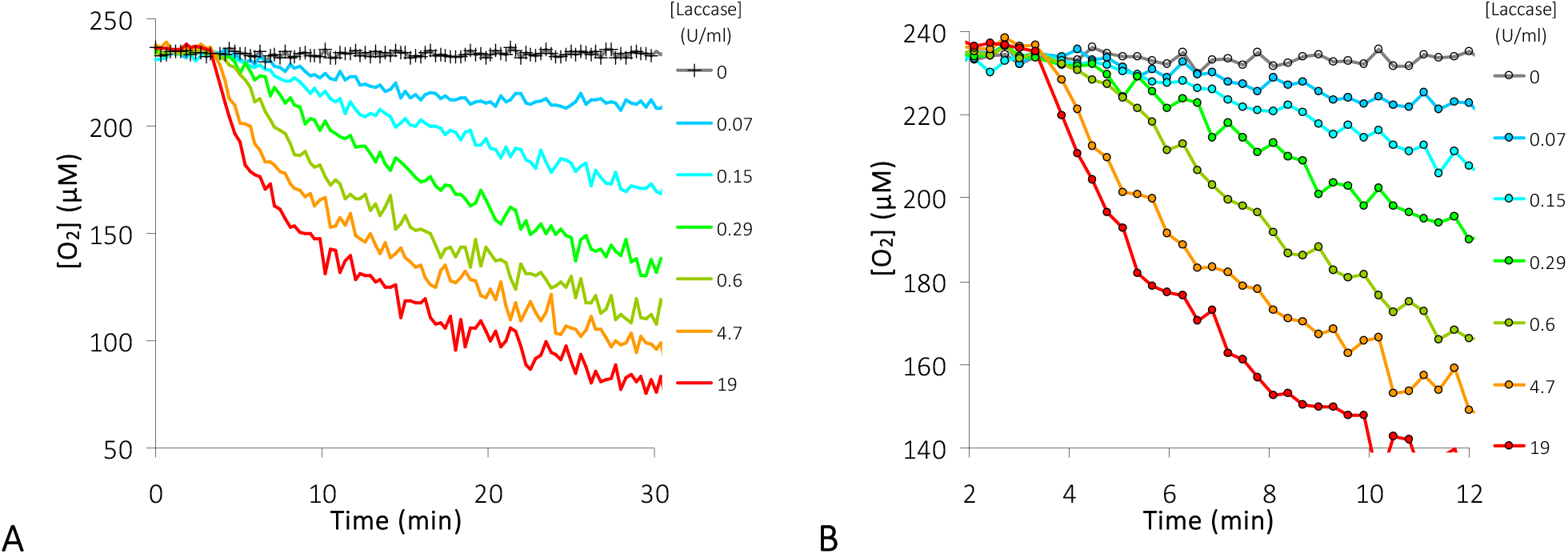
Consumption of O_2_ by different laccase activities recorded ratiometrically using Pd-Benzoporphyrin fluorescence. **A:** The reaction was started at t = 3.3 min by injection of pyrogallol into a reaction milieu and wells were immediately sealed. **B:** Enlargement of a data section from **A**. Final assay conditions: CPB pH=5.5, [Pyrogallol] = 6 mM; 28.2°C; Salinity 5.1 mS/cm; F _Sal_ = 0.984; [Laccase] as indicated in the inset legends; Sample rate = 1/18·s^-1^.

### O_2_ consumption by respiration during bacterial growth

Growth of, and fermentation by aerob microorganisms in batch cultures is restricted by nutrients and O_2_ supply. The effect of oxygen deficiency on bacterial growth can be monitored with optical oxygen sensors during the whole culture period, when the optical density (OD) of the culture is recorded in parallel.

*Vibrio natriegens* is one of the fastest growing bacterial species. It can reach doubling times as low as 10 min under aerob conditions^56^. Here *V. natriegens* was grown in clear MTPs coated with NMP in order to monitor cellular density (OD_600_) and [O_2_] simultaneously (**Fig. 10**). With sealed wells, exponential growth started after a lag phase of ca. 2.6 h and O_2_ consumption started in parallel. Growth levelled off when [O_2_] dropped below 20 µM. Removal of the sealing foil, to allow for O_2_ ingress, caused further growth by a factor of 1.6 (OD_600_ increase from 0.8 to 1.3), indicating that growth retardation within the first 20 hours was mainly restricted by the lack of gas exchange. The second growth period stopped at OD_600_ = 1.3 (ln(OD_600_)=0.26), suggesting that nutrients were exhausted. The O_2_ ingress seen during the last 12 h of the experiment (**Fig. 10**, blue line) reflects the fact that respiration of the cells ceased.

**Figure 10.**
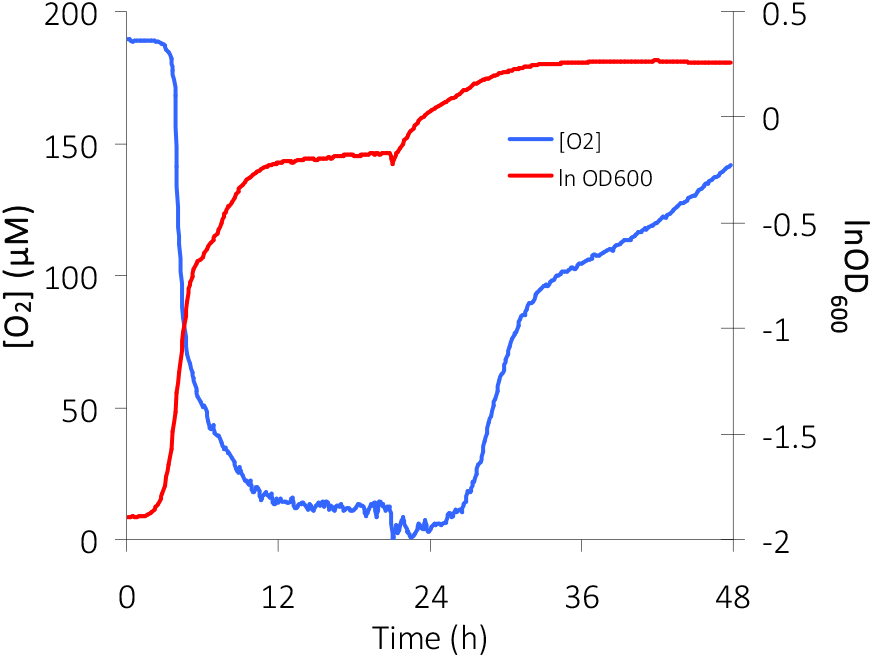
Growth and respiration of *Vibrio natriegens* monitored in MTP wells. The [O_2_] measured by NMP fluorescence ratiometry (left ordinate, blue line) was recorded in parallel to OD_600_ (right ordinate, logarithmic scale, red line). The maximum growth rate coincides with maximum respiration at t = 4.3 h. The sealing foil was removed at t = 21 h, allowing O_2_ ingress. This resulted in further growth, whereas growth retardation (from t = 27 h onwards) leads to less O_2_ consumption and consequently increased O_2_ ingress. Culture conditions: LB+v2-medium; 28°C; salinity 44 mS/cm; F_Sal_ = 0.85; linear shaking 1.5 mm; AVR of n = 33; SD < 12%; sample rate = 1/5 min^-1^

It appears reasonable to assume that the O_2_ shortage is the cause for growth retardation when MTP wells are sealed and that growth resumes as soon as gas exchange is restored, provided nutritional resources are still available. However, there may be other decisive factors, such as pH and bicarbonate, which are affected by bacterial growth when gas exchange is blocked. If needed, pH could be also monitored by fluorescence ratiometry in MTPs using specific immobilized pH-sensitive fluorescein derivatives ^21^.

### O_2_ production by the catalase reaction

Catalase catalyses the disproportionation of hydrogen peroxide (H_2_O_2_) (**Eq. 7**) and in consequence produces O_2_ (**Fig. 11**). The O_2_ increase is efficient and quick when H_2_O_2_ and catalase are combined (**Fig. 11 B**). This allows to estimate the response time of the embedded O_2_-indicator relative to the used sample rate.

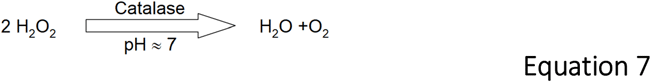

**Figure 11.**
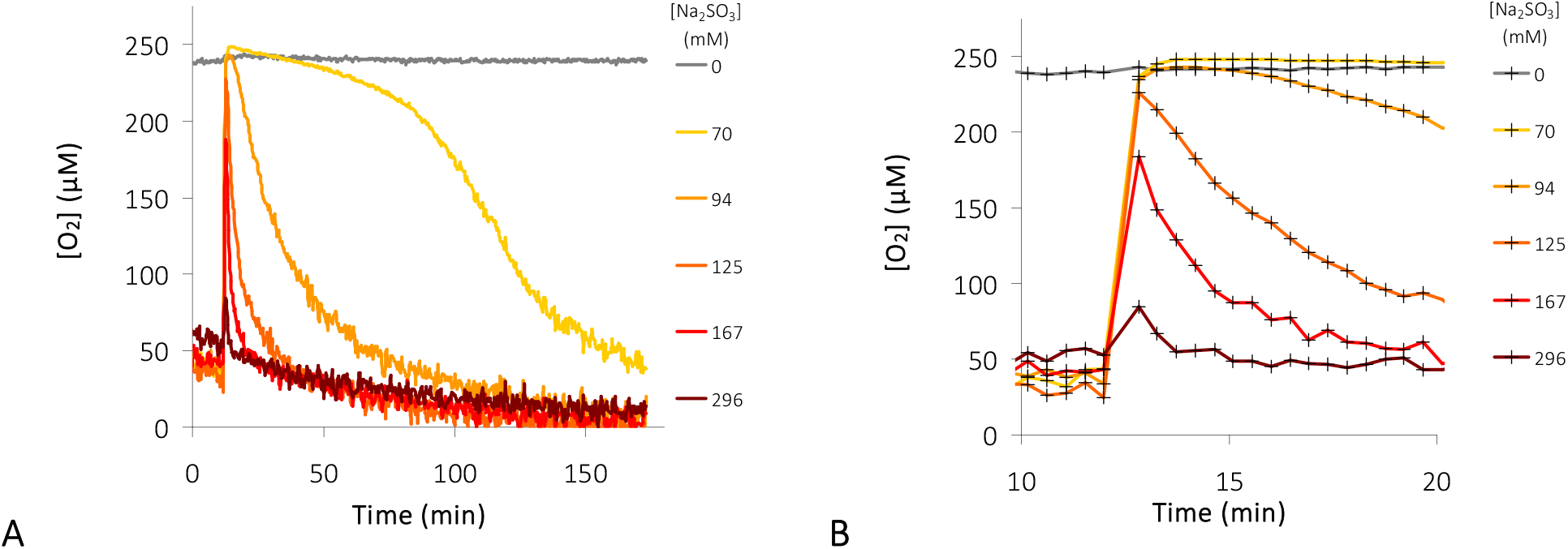
O_2_ bursts produced by catalase. monitored in presence of different sulfite concentrations. **A** : The lower [Na_2_SO_3_], the more pronounced appears the catalase induced O_2_ burst. The reactions were started at t = 12 min by injection of H_2_O_2_ into a sulfite solution containing catalase. [O_2_] was recorded ratiometrically with Pd-Benzoporphyrin fluorescent indicator. **B:** The close-up indicates an instantaneous response of the indicator. Final assay composition: KPB 50 mM pH = 7; 28°C; [H_2_O_2_] = 21 mM; [Na_2_SO_3_] indicated by inset legends; salinity (F_Sal_; **Eq. 11**) taken into account. Sample rate ca. 2·min^-1^.

The experiment (**Fig. 11**) demonstrates that the O_2_ bursts released by the catalase reaction (**Eq. 7**) is compensated (‘buffered’) by Na_2_SO_3_ (**Eq. 5**) as long as [Na_2_SO_3_] is high enough. Thus, with high [Na_2_SO_3_], the free [O_2_] in the medium remains largely unaffected by the catalase reaction (dark red line). However, with lower [Na_2_SO_3_] the released O_2_ causes noticeable perturbations of [O_2_] in the medium because the compensation by Na_2_SO_3_ is slower than production by catalase. The indicator shows a rapid response relative to the data sampling frequency and immediately displays the change in [O_2_] without delay. From this, it can be concluded that the coating is thin enough (thickness of indicator film ≈ 1.5 µm) to ensure quick response, although the diffusion of O_2_ is more than 100 times slower in the polystyrene film (D = 1·10^-7^ cm^2^·s^-1^; ^57^) when compared with water (D ≈ 2·10^-4^ cm^2^·s^-1^; ^46,58^).

### Photosynthetic O_2_ production and respiration by unicellular algae

Photosynthetic organisms produce O_2_ in excess when the intensity of photosynthetic active radiation (PAR) is above the light compensation point. A sufficient dose of light leads to an [O_2_] increase in the microalgae suspension, whereas darkness allows monitoring the respiration. This offers a variety of experiments to answer diverse questions about light compensation point, the interaction of photosynthesis and respiration and the effect of specific inhibitors.

The data displayed in **Fig. 12** show that the dose of light (i.e. amount of photons applied) determines whether a balance between photosynthesis and respiration is established or whether an O_2_ deficiency, approaching anoxia, occurs.

**Figure 12.**
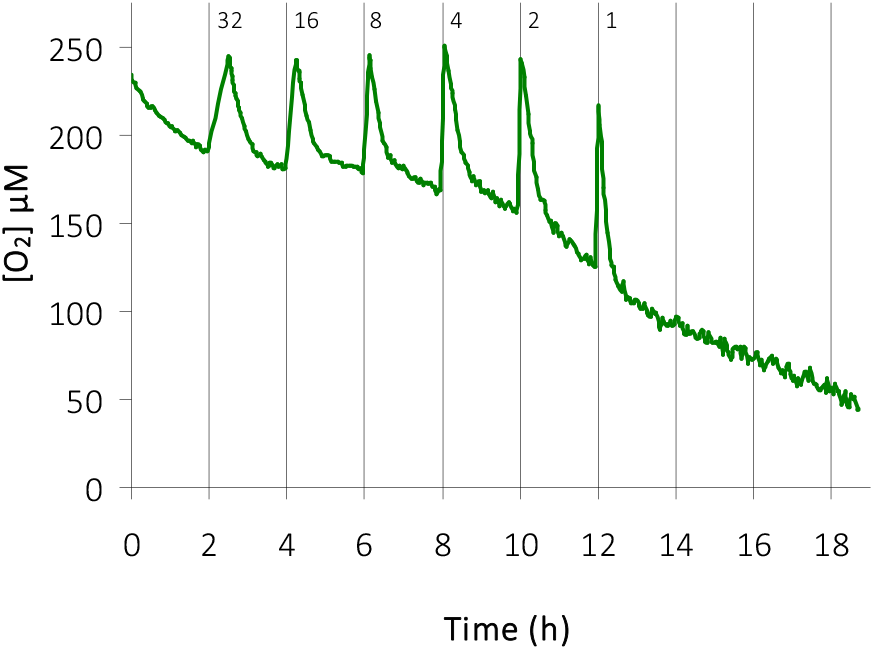
O_2_ consumption and production by microalgae. (*Eremosphaera viridis*) monitored using Pd-Benzoporphyrin fluorescence in sealed MTPs. Respiration of algae was recorded for several hours. Every 2 hours, the cells were irradiated with PAR (100 µmol·s^-1^·m^-2^) for several minutes. Numbers above each peak indicate the duration of irradiation in minutes. Variation of the light periods demonstrate that respiration cannot be compensated by less than 4 min of light. Experimental conditions: 200 µl algal suspension with ca. 14,000 cells per well; ¼ M & S culture medium; 27°C, 1.2 mS/cm; no salinity correction with calibration. Sample rate = 1/3·min^-1^.

**Fig. 13** demonstrates the effect of 3-(3,4-dichlorophenyl)-1,1-dimethylurea (DCMU), a specific inhibitor of photosynthesis ^59–61^. The evolution of O_2_ during light periods is inhibited by DCMU (**Fig. 13**; yellow trace). Since DCMU is soluble only in Ethanol (EtOH), a control without DCMU was run im parallel, showing that EtOH has a pronounced positive effect on respiration (**Fig. 13**; blue trace) when compared with the control (green trace). This latter effect is in line with earlier findings, showing that ethanol can serve as carbon source for microalgae and stimulates metabolism and biomass production ^62,63^.

**Figure 13.**
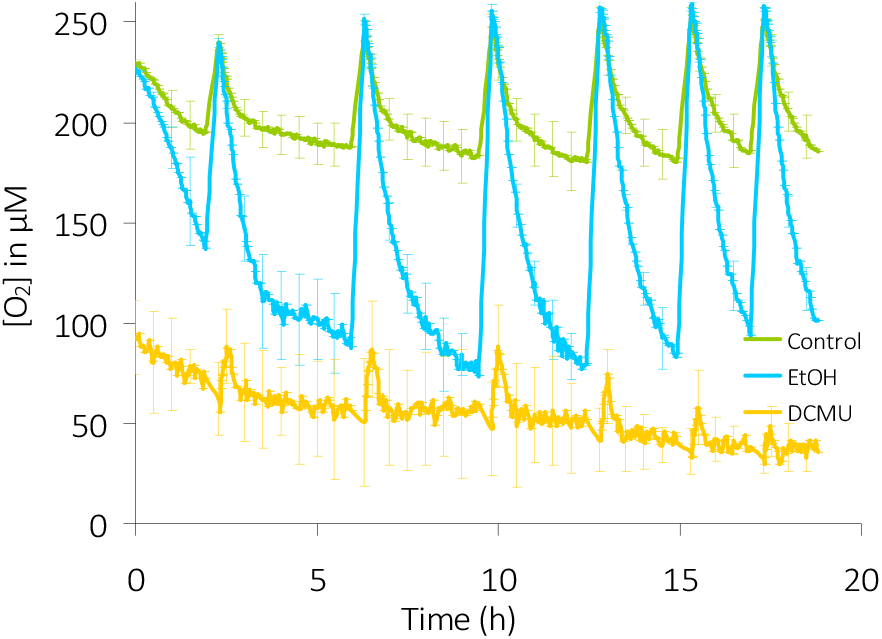
Effect of DCMU and ethanol on photosynthesis and respiration of microalgae. (*Eremosphaera viridis*) Wells coated with Pd-Benzoporphyrin were filled with 200 µl of algal suspension (7000 cell/ml) and sealed with adhesive transparent polyester-foil. [O_2_] was monitored in absence of any inhibitor (green curve; ‘Control’), in presence of Ethanol (0.5%; blue curve), and in presence of DCMU (100 µM) together with Ethanol (yellow curve). Algae were irradiated at predetermined intervals with PAR of 25 µmol·s^-1^·m^-2^ for 20 minutes. AVR of n = 12; error bars represent SD. Experimental conditions: ¼ M & S culture medium; 1.2 mS/cm; no salinity correction with calibration; 26°C; Sample rate = 1/3·min^-1^.

The experiment (**Fig. 13**) has been repeated with diverse concentrations of DCMU and EtOH to roughly estimate the effective minimal concentrations needed for the effects seen (Supporting information: **Fig. SI 5** - Effect of DCMU and ethanol on photosynthesis and respiration of algal cells). A pronounced inhibition of photosynthetic O_2_ evolution can be seen with [DCMU] > 10 µM (**Fig. SI 5 B**; yellow curve). Enhanced respiration, is seen already with 0.03% EtOH (= 6.8 mM) and increases further with increasing [EtOH] (**Fig. SI 5 A** to **F**; blue curves).

### Oxygen ingress needs to be kept in check

O_2_ ingress and egress are undesirable effects when monitoring [O_2_] in MTPs, because the recorded fluorescence should reflect only those [O_2_] changes, which are attributable to the biochemical reactions in the MTP wells. Hence, these effects have to be kept in check. This is demonstrated in the supporting information (**Figs. SI 7 – SI 10**). O_2_ ingress could be minimized to a negligible level by sealing the reaction volume from the atmosphere. Another strategy is to monitor these effects separately, in order to eliminate the effect when processing the experimental data. A procedure, how to quantify oxygen ingress has been demonstrated above (**Fig. 7**) and various measures have been proposed to minimise these effects ^34^. Among them was the recommendation to overlay the reaction volume with oil. However, even oil is permeable for O_2_ and ingress is only slowed down. This is demonstrated in the supporting information **Figs. SI 6, SI 7**. Wells containing solutions with different amounts of Na_2_SO_3_ were covered with different volumes of oil and O_2_ ingress was monitored. The data show that O_2_ ingress can be reduced by a reasonable layer of oil, but cannot be entirely prevented. This is in line with results presented by Arain et al. 2005 ^34^.

When sealing the MTP with adhesive film, a residual air volume with 21 % O_2_ remains over the liquid volume to be probed in the well. This still causes unwanted O_2_ ingress as demonstrated in the supporting information **Fig. SI 8**. However, the data show that minimising the residual air volume in the wells reduces the O_2_ ingress considerably. There are diverse adhesive films and foils on the market developed specifically to cover MTPs. We tested a few for their suitability to reduce or stop unwanted O_2_ ingress and found, that polyester cover film initially developed for sealing PCR reactions is optimal. The film is transparent, allowing simultaneous OD_600_ and [O_2_] measurements (**Fig. 10**), is largely impermeable to O_2_, and thus reduces O_2_ ingress to a negligible degree when the residual entrapped air volume is small.

### O_2_ binding capacity of plastic material

Polystyrene (PS) is known for its O_2_ permeability ^34^. O_2_ quickly diffuses into PS and quenches the fluorescence of embedded porphyrins (**Fig. 11**). Despite the low diffusion coefficient of O_2_ in PS, the effect is quick since the indicator layer is thin (i.e. ≈ 1.5 µm). However, the whole MTP is made of PS and thus soaks up O_2_ like a sponge. Due to the absorption capacity of PS for molecular oxygen ^44^, bound O_2_ can diffuse out of the plastic material into the reaction volume, where [O_2_] is to be monitored ^64^. This effect is seen in the data demonstrated by **Fig. SI 10** in the supporting information.

Wells of MTPs incubated overnight in an O_2_-free atmosphere in presence of O_2_-absorber were filled with Na_2_SO_3_ solution and fluorescence ratios were recorded for several hours. Two different effects can be seen (**Fig. SI 10**, red dots). First, the O_2_ is consumed by the sulfite oxidation reaction (**Eq. 5**) and fluorescence ratio increases within the first 40 min due to the [O_2_] decrease. Second, this process of O_2_ consumption is superseded by the diffusion of O_2_ from the ambient atmosphere into and through the O_2_-free MTP plastic material, leading to a ratio decrease and finally to an equilibrium at around R = 5. Control data (**Fig. SI 10** blue dots) show, that O_2_ diffusing through and released from the O_2_-saturated plastic is largely consumed by the sulfite (**Eq. 5**), and the same equilibrium value is finally reached.

In summary, O_2_ ingress cannot be completely avoided. However, there are means to minimise this interfering effect. An important factor is the timing of the experiment. It is possible to estimate whether diffusion and ingress are negligible in comparison to the effect to be quantified (e.g. respiration or photosynthesis - **Figs. 12, 13**). Finally, appropriate controls run in parallel will allow for a correction of the experimental data.

### pH-dependence of NMPs entrapped in PS

The NMP fluorescence is independent from pH of the samples in the wells, because the indicator is trapped in the plastic material (PS) and thus changes of [H^+^] should not have an impact. This statement does not hold for any liquids in the wells because PS, when sulfonated, becomes proton conductive ^65–67^.

To demonstrate this effect, MTP wells doped with **1** or **2** were filled with buffers of different pH and fluorescence was measured. Sulfonic acid buffers cause a significant pH dependence, revealing a H^+^-permeability of PS (**Supporting information: Figs. SI 11A, B** and **SI 12A, B**; pH-dependence of NMPs embedded in PS). Mineral buffers, in contrast, do not show this effect (**Figs. SI 11C, D** and **SI 12C, D**). Hence, sulfonic buffer components in the reaction volume, such as MES, HEPES, and TAPS, make PS proton permeable and will affect indicator fluorescence.

### Advantages

Well-to-well variations of fluorescence emission resulting from inhomogeneities in the indicator coating are eliminated by ratioing fluorescence intensities taken at two different wavelengths. The Pd-Porphyrins **1** and **2** introduced here (**Fig. 1**) are inherent ratioable indicators (**Fig. 5**). In contrast to previous studies, no additional O_2_-insensitive reference dye is needed to obtain ratioable fluorescence emissions. With Pt-Porphyrins and a co-immobilized O_2_-insensitive reference dye (e.g. Rhodamine) a dynamic signal response factor of R_max_/R_min_ < 3 can be achieved ^21,38,39^. However, with Pd-Porphyrins, which require no reference dyes, larger response factors are possible.

The indicator layers obtained with the coating protocol proposed here are clear and not turbid (**Fig. SI 1**). This bears the advantage that absorbance or optical density (OD) can be quantified simultaneously with [O_2_] in each well (**Fig. 10**). The coating procedure is simple (see experimental section) and quick. Coating of 10 MTPs can be completed within 45 min and the costs per plate are below 5 € per plate (material only, labour not included).

## Conclusions

Pd-Porphyrins are superior O_2_-indicators for use in MTPs to monitor the concentration of oxygen dissolved in aqueous solutions. The novel coating protocol provided here allows an inexpensive, fast, and easy production of MTPs for ratiometric optical readout. The conversion of fluorescence intensity ratios in terms of oxygen concentrations is straightforward. Thereby, temperature and salinity effects can be taken into account. In summary, the MTPs prepared with immobilised NMP, as proposed with this study, meet all quality standards for spectroscopic sensors ^68^. The methods reported here open up a wide field of diverse research approaches, provided critical frame parameters are known and suitable control experiments are carried out.

## Experimental Section

### Specific Instrumentation and Materials

Plate reader Infinite M200 PRO with an injection unit (Tecan, Crailsheim, Germany) was used for optical readouts such as absorbance, optical density and fluorescence (bottom read). Suitable scripts were programmed via the menu interface of the manufacturer’s i-control™ software.

Specific Chemicals Chloroform;

### Petroleum-Benzine 40-60

Palladium (II) 5,10,15,20-(tetrapentafluorophenyl) - porphyrin (CAS 72076-09-6;MW = 1079 g/mol) Palladium (II) 5,10,15,20-(tetraphenyl) - tetrabenzoporphyrin (CAS 119654-64-7; MW = 919 g/mol) Sources and all other standard chemicals used are listed in the Supporting Information (**Tab. SI 1**).

### Specific Consumables and Specific Disposable Material

Standard clear flat bottom 96-Well PS-MTP Sarstedt #82.1581

Aluminiumpaper sandwich film Carl Roth # X172.1

Transparent Polyester sealing film Carl Roth # APT8.1

Standard consumables such as tubes, pipette tips and dishes were from Sarstedt, Roth, or Greiner.

### Living Material

*Eremosphaera viridis* (De Bary)

*Vibrio natriegens* (Vmax)

### Buffers, Culture Media, Stock Solutions, and Coating Solutions

CPB (Citric acid Phosphate Buffer) 50 mM Citric acid adjusted with K_2_HPO_4_ to pH = 5.5

KPB (Potassium Phosphate Buffer) 50 mM K_2_HPO_4_ adjusted with 50 mM KH_2_PO_4_ to pH = 7.4

#### Algal culture medium (¼ M&S)

¼ Murashige & Skoog Medium: 1 g MS-salts plus 1 g MES dissolved in 1 L dH_2_O adjusted with KOH to pH = 6.2

#### Bacterial culture medium (LBv2)

LB-medium supplemented with v2 salts (i.e. 204 mM NaCl, 4.2 mM KCl, and 23.14 mM MgCl_2_).

#### PS-stock solution 5 % w/v in Chloroform

5 g Polystyrene (from petridishes) dissolved in 100 ml Chloroform

#### Chloroform Petrol-Benzine Mix (CB-Mix 1 : 1)

100 ml Petroleum-Benzine mixed with 100 ml Chloroform

Fluoroporphyrine stock solution: 1 mg Pd-Fluorophorphyrin (**1**; MW = 1079 g/mol) dissolved in 400 µl PS-stock gives a deep purple-red colour with about 2.3 mM of **1**.

Benzoporphyrine stock solution: 1 mg Pd-Benzophorphyrin (**2**; MW = 919 g/mol) dissolved in 800 µl PS-stock gives a rich green colour with about 1.4 mM of **2**.

Fluorophorphyrine coating solution: 400 µl of Fluorophorphyrine stock solution mixed with 11.6 ml of CB-Mix. The coating solution contains 51,7% Chloroform (v/v), 48,3% Petrol-Benzine (v/v), 0.17% Polystyrene (w/v), and 83.3µg/ml Fluorophorphyrine (= 77 µM)

Benzophorphyrine coating solution: 800 µl of Benzophorphyrine stock solution mixed with 23.2 ml of CB-Mix. The coating solution contains 51,7% Chloroform (v/v), 48,3% Petrol-Benzine (v/v), 0.17% Polystyrene (w/v), and 41.6µg/ml Benzoporphyrine (= 45 µM)

## Methods

### Coating Procedure

The coating is an improved and largely simplified procedure originally adapted from Borisov et al. ^19^ and De Moraes Filho et al. ^42^.

From the coating solution, 30 µl are dispensed in each well of flat-bottom MTPs and solvent vapours are evaporated overnight in a fume hood. This gives a homogeneous clear transparent coating as shown in the supporting information (**Fig. SI 1**), with ca. 1 to 3 µg per well of NMP embedded in a PS layer of about 1.5 µm thickness.

Note: Coating solutions also depends on the type of MTP to be coated. Flat bottom MTPs were preferably used here since they allow recording of OD_600_ and fluorescence in parallel. For U-bottom MTPs, the coating solution should have the threefold concentration of porphyrin (i.e. 400 µl Fluorophorphyrine stock on 3.6 ml CB-Mix, or 800 µl Benzophorphyrin stock on 7.2 ml CB mix). Only 10 µl of the concentrated coating solution are needed to be dispensed per U-well for a sufficiently homogeneous clear coating.

### Fluorescence Recording

Wells of coated plates were loaded as described in the results section. Fluorescence and absorption readouts were recorded with the above mentioned plate reader using specifically programmed i-control™ scripts.

### Calibration of NMP fluorescence ratios in terms of [O_2_] at constant temperature and zero salinity

A reasonable first approach for converting fluorescence ratios R into oxygen concentrations [O_2_] is the assumption of a strict linear relationship (**Eq. 8**), provided the maximum O_2_ solubility [O_2_]_max_ is known together with the given salinity and temperature of the probed solution and the atmospheric pressure. R_max_ is the maximum fluorescence ratio obtained under anoxic conditions ([O_2_] = 0 µM), whereas R_min_ is the ratio at maximum dissolved O_2_ in aqueous solution ([O_2_] = [O_2_]_max_).

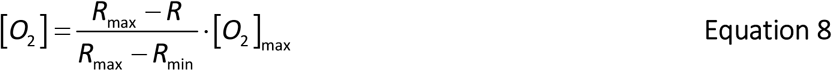

According to DOTABLES^3^ (**Supporting information: Fig. SI 2**), [O_2_]_max_ is around 280 µM at RT = 20°C, in fresh water (i.e. no salinity) and with normal atmospheric pressure of around 100 kPa.

### Accounting for temperature effects when calibrating NMP fluorescence ratios in terms of [O_2_]

With increasing temperature θ (°C), the equilibrium O_2_ solubility (i.e. [O_2_]_max_) decreases in aqueous solutions ^2,69^. To account for this effect, the above calibration equation (**Eq. 8**) needs to be extended with a temperature-dependent term. Therefore, exponentials were fitted to data from dissolved oxygen tables (DOTABLES) ^3^ to obtain continuous model functions describing the dependence of [O_2_]_max_ on temperature θ (**Eq. 9; supplemental Information Fig. SI 2**).

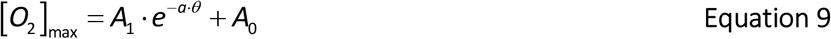

This term (**Eq. 9**) is used to correct for temperature effects, when experiments were performed under varying temperatures. The resulting extended calibration equation is (**Eq. 10**):

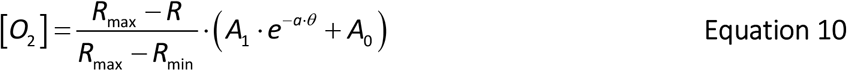

For instance, the parameter triple of the model function (Eq. 9) fitted to the 100 kPa data set (**Fig. SI 2** red dots) is (A_1_; a; A_0_) = (323; 0.037; 126). The necessity of temperature correction is discussed and demonstrated in the results section (**Fig. 8**).

### Considering salinity with calibration

In addition to the temperature dependence, there is also a salinity dependence of oxygen solubility in aqueous media ^70^. [O_2_]_max_ decreases with increasing ion concentrations, which are typically expressed in units of specific conductivity (S/cm) of the solution. For more precise calibration, recorded [O_2_] needs to be corrected by a correction factor F_Sal_ ^2^ to take account for the effect of salinity (**Eq. 11**).

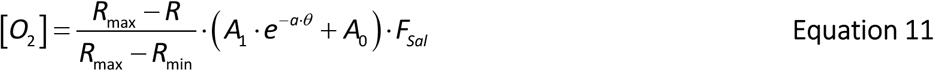

Values of F_sal_ for a series of salinities are shown in the supporting information (**Fig. SI 3**) and can be obtained online from DOTABLES ^3^.

A barometric correction was omitted, since all data presented in the context of this study were obtained at sea level and weather-related changes in atmospheric pressure were neglected.

For experiments requiring calibration, two types of references were established:

1. Wells were filled with 30 mg O_2_-absorber powder (i.e. iron charcoal salt mix) each and sealed with adhesive polyester-foil to obtain maximal fluorescence and R_max_ at [O_2_] = 0.
2. For saturation, i.e. to obtain minimal fluorescence and R_min_ with [O_2_] = [O_2_]_max_, wells were filled with distilled water and left open for O_2_ exchange with the ambient air.

## Associated Content – Supporting Information (SI)

**Appearance of transparent MTPs coated with Pd-porphyrin in PS**

Figure SI 1 Images: Flat-bottom MTPs for [O_2_] quantification

**Temperature dependence of oxygen solubility in aqueous solutions**

Figure SI 2 The equilibrium solubility of atmospheric molecular oxygen in water.

**Salinity dependence of oxygen solubility in aqueous solutions**

Figure SI 3 The salinity correction factor F_Sal_

**The linear relationship between fluorescence ratios and [O**_**2**_]

Figure SI 4 Correlating NMP fluorescence ratios with amperometric oxygen recording

**The Effect of DCMU and Ethanol on Photosynthesis and Respiration**

Figure SI 5 Effect of DCMU and ethanol on photosynthesis and respiration of algal cells.

**The O**_**2**_ **ingress from ambient air**

Figure SI 6 O_2_ ingress reduced with oil. Figure SI 7 The effect of oil on the O_2_ ingress

**O**_**2**_ **ingress from residual air volume entrapped under the sealing film**

Figure SI 8 The dependence of O_2_ consumption on residual entrapped air volumes.

Figure SI 9 O_2_ ingress observed with different entrapped air volumes

**O**_**2**_**-diffusion into and through polystyrene**.

Figure SI 10 O_2_-absorption by and diffusion through the MTP plastic material.

**pH-dependence of NMPs embedded in polystyrene**

Figure SI 11 The pH-dependence of Pd-Fluoroporphyrin fluorescence emission

Figure SI 12 The pH-dependence of Pd-Benzophorphyrin fluorescence emission.

## Authors’ contributions

C.P. conceived the study, evaluated data and wrote the first draft of the manuscript. D.P. carried out the experiments reported in the main manuscript, performed data processing and participated in amending the draft. Both authors approved the final version.

## Notes and Declaration

Both authors declare that they have no conflicts of interest.

## Acknowledgements

We thank Benoit Habermeyer of PorphyChem SAS for recommending **1** and **2** as appropriate O_2_-sensor molecules. The authors are grateful to Isabel Moreau (BiMo; Kiel University) for inspiring tips, to Axel Scheidig (Structural Biology Group, Kiel) for his generous support, and to Sonja Vollbehr (BiMo; Kiel University) for technical assistance. BBE Moldaenke (Schwentinental) and WTSH (Kiel) generously provided the plate reader. Finally, we gratefully acknowledge access to the core facilities of the Zentrum für Biochemie und Molekularbiologie (BiMo), Christian-Albrechts-Universität, Kiel.

## Abbreviations

1: Palladium(II)-5,10,15,20-(tetrapentafluorophenyl)porphyrin (CAS 72076-09-6)
2: Palladium(II)-5,10,15,20-(tetraphenyl)tetrabenzoporphyrin (CAS 119654-64-7)
[M]: a molecular species M in brackets designates the concentration in the solution given
Avr: average
a.U.: arbitrary units
DCMU: 3-(3,4-dichlorophenyl)-1,1-dimethylurea
EtOH: Ethanol
FWHM: full width at half maximum
HTP: high throuput
KPB: potassium phosphate buffer
MTP: microtiterplate
MW: molecular weight
OD: optical density
NMP: noble metal porphyrin
PAR: photosynthetic active radiation
PS: polystyrene
SD: standard deviation
SI: supporting information
S-PS: sulfonated polystyrene

## Variable Declaration

τ (*greek letter: tau*): time constant
θ (*greek letter: theta*): temperature
F: fluorescence
R: Ratio of fluorescence intensities
T: integration time
t: time

