## Supporting Information for "Ratiometric Quantification of Dissolved Molecular Oxygen in Microplates for Biochemical Assays Using Palladium Porphyrin Photoluminescence"

### Appearance of transparent MTPs coated with Pd-porphyrin in PS

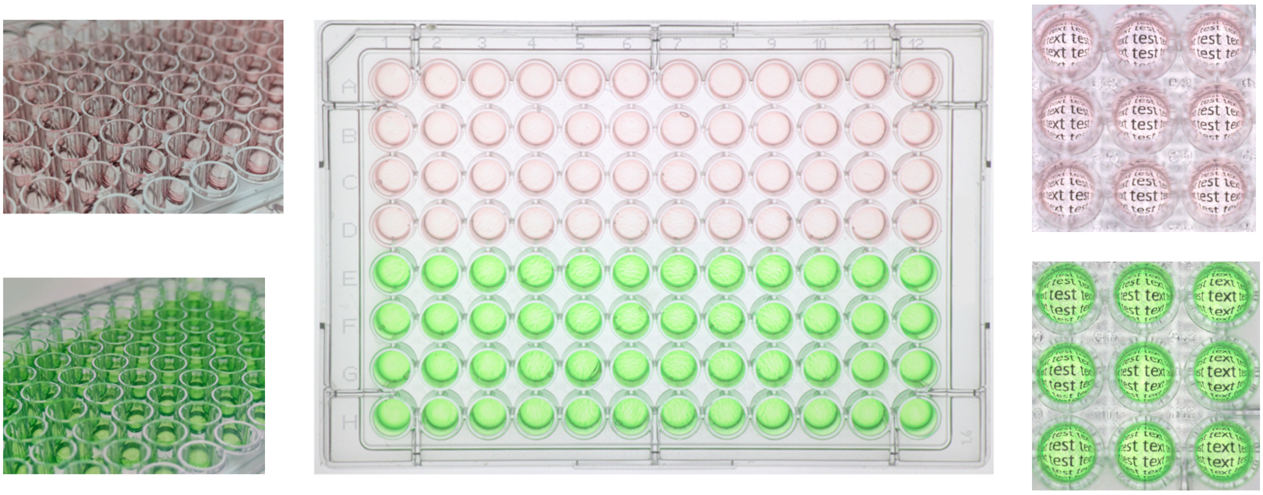

**Figure SI 1 Flat-bottom MTPs for [$O_2$] quantification**. Wells coated with **1** (pink) and **2** (green) as described in the experimental section. The close-ups on the right hand side demonstrate clear transparent coating.

### Temperature dependence of oxygen solubility in aqueous solutions

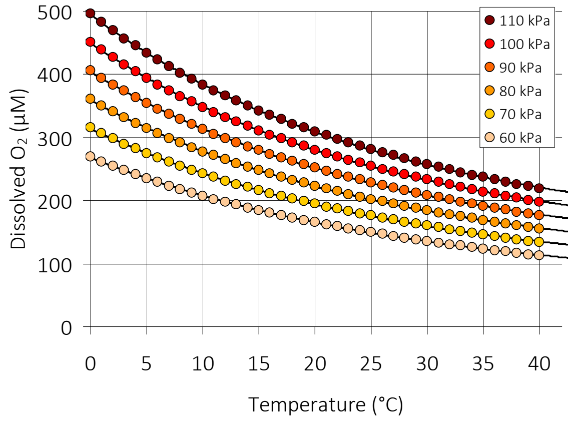


**Figure SI 2 The equilibrium solubility of atmospheric molecular oxygen in water.** Presented are the data from DOTABLES[1], which originate from Benson and Krause 1984 [2]. Given is the dependence of the maximum $O_2$ concentration in freshwater (i.e. $[O_2]_{max}$ on y-axis) on temperature (x-axis) and on barometric pressure of air with 21% $O_2$ (inset legend). Single exponential model functions model functions

$$[O_2]_{max} = A_1 \cdot e^{-a \cdot \theta} + A_0$$

were fitted to the data-sets for each pressure value, describing the dependence of $[O_2]_{max}$ on temperature $\theta$. At 20°C and normal barometric pressure near sea level (≈ 100 kPa) the equilibrium solubility of $O_2$ in water is about 280 µM.

### Salinity dependence of oxygen solubility in aqueous solutions

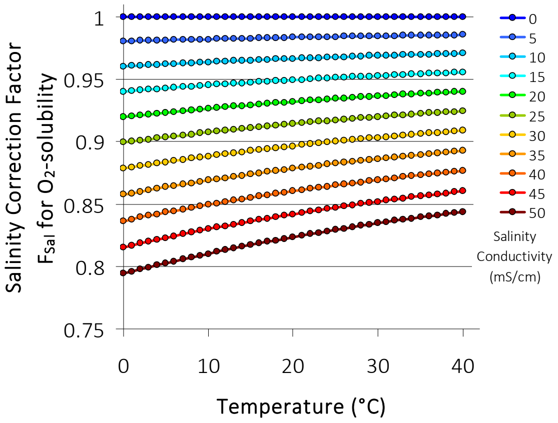


**Figure SI 3 The salinity correction factor $F_{Sal}$**
The solubility of $O_2$ depends on the salinity of the aqueous solution (i.e. ion concentration). Ion concentrations are expressed in units of specific conductivity (mS/cm) of the solution. For a more precise calibration, $[O_2]$ values obtained optically need to be corrected by a correction factor $F_{Sal}$ to take account for the effect of salinity (**Eq. 11** of the main text). Given is the correction factor $F_{Sal}$ (y-axis) in dependence on temperature (x-axis) for diverse degrees of salinity (inset legend). Data are based on Benson and Krause 1984 [2] and are taken from DOTABLES [1].

### The linear relationship between fluorescence ratios and $[O_2]$

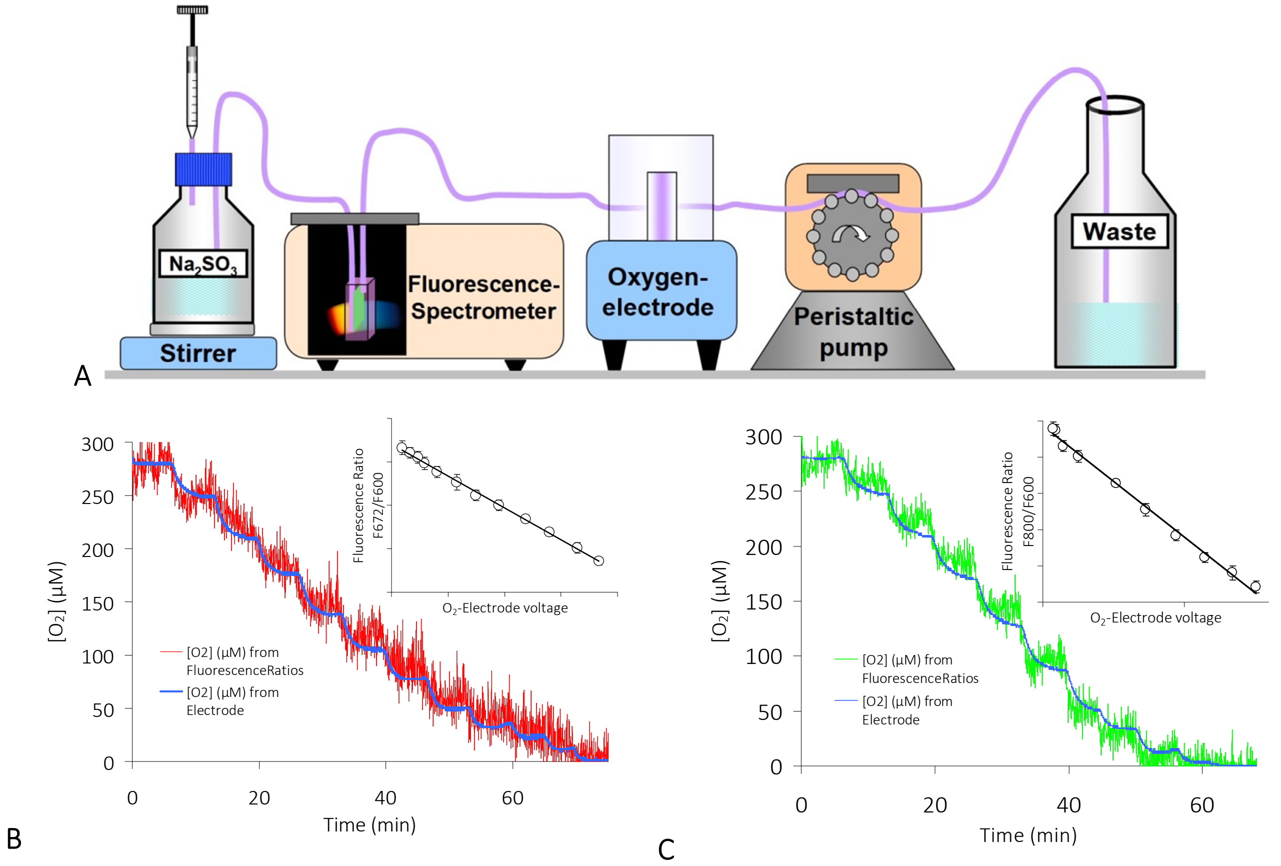


**Figure SI 4 Correlating NMP fluorescence ratios with amperometric oxygen recording**. A: A piece of NMP-coated PS, mounted diagonally in a flow-through cuvette, was placed in a fluorescence spectrometer (Varian Cary eclipse) and connected in series with a flow-through Clark-type $O_2$-electrode (Rank Brothers Ltd. Model 10). The set-up was perfused with water. $[O_2]$ was adjusted by stepwise addition of $Na_2SO_3$ to the storage volume (bottle left hand side). Fluorescence of immobilised **1** (B) and **2** (C) in the cuvette and the electrical current from the $O_2$-electrode were recorded in parallel. Sample rate = $1 \cdot s^{-1}$. **Insets**: The $O_2$ dependent fluorescence emission ratios (i.e. F672/F600@$F_{ex}$554 for **1** and F800/F600@$F_{ex}$445 for **2**). The amperometric signals exhibit a linear relationship with $R^2 = 0.998$ for **1** (B) and $R^2 = 0.994$ for **2** (C), revealing that fluorescence quenching by $O_2$ is predominantly dynamic and thus a Stern-Volmer process. AVR of n ≥ 40; error bars represent SD.

### The Effect of DCMU and Ethanol on Photosynthesis and Respiration

The effect of 3-(3,4-dichlorophenyl)-1,1-dimethylurea (DCMU), a specific inhibitor of photosynthesis [3–5], on photosynthetic $O_2$ evolution by unicellular algae was investigated. Since DCMU needs to be dissolved in ethanol, appropriate controls with ethanol lacking DCMU were run in parallel.

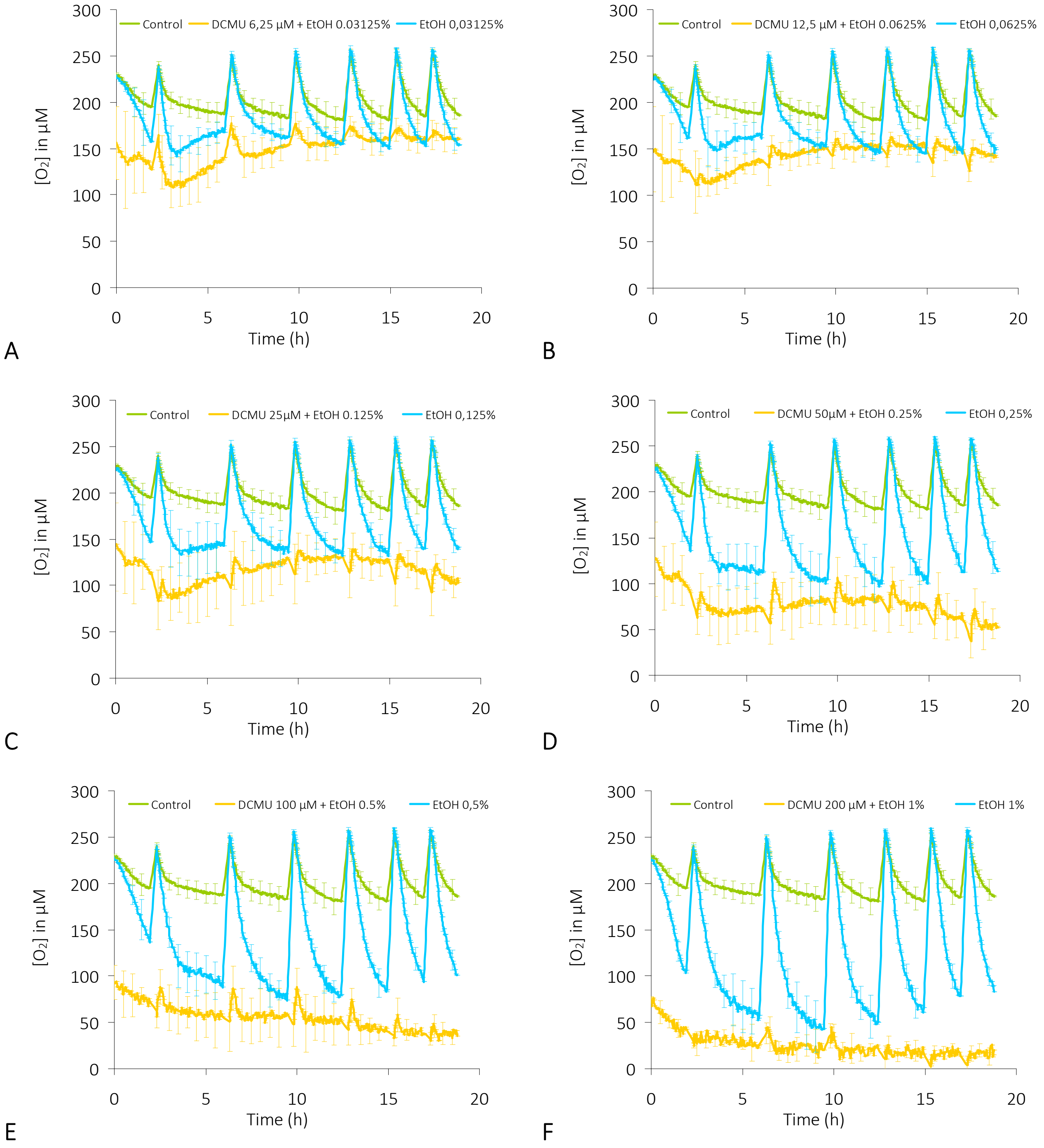


**Figure SI 5 Effect of DCMU and ethanol on photosynthesis and respiration of algal cells** Wells coated with Pd-Benzoporphyrin were filled with 200 µl of algal cell suspension *(Eremosphaera viridis;* ≈ 7000 cell/ml) and sealed with adhesive transparent polyester-foil. $[O_2]$ was monitored in absence of any effector (green curves; 'Control'), in presence of ethanol (blue curves), and in presence of DCMU together with ethanol (yellow curve). Algae were irradiated at predetermined intervals with PAR of 25 $\mu mol \cdot s^{-1} \cdot m^{-2}$ for 20 minutes. Concentrations [DCMU] and [EtOH] increasing from A to F are given in the inset legends of each panel. AVR of n = 12; error bars represent SD. Experimental conditions: ¼ M & S culture medium; conductivity 1.2 mS/cm; no salinity correction with calibration; 26°C; sample rate = $1/3 \cdot min^{-1}$.

### The $O_2$ ingress from ambient air

Diverse techniques have been suggested to minimise oxygen ingress from the atmosphere when monitoring $[O_2]$ in MTPs. In particular, the overlay of oil onto the aqueous volume in the wells has been recommended [6]. The following data demonstrate how oxygen ingress is reduced by adding oil to the aqueous solution in the wells (**Fig. SI 6**).

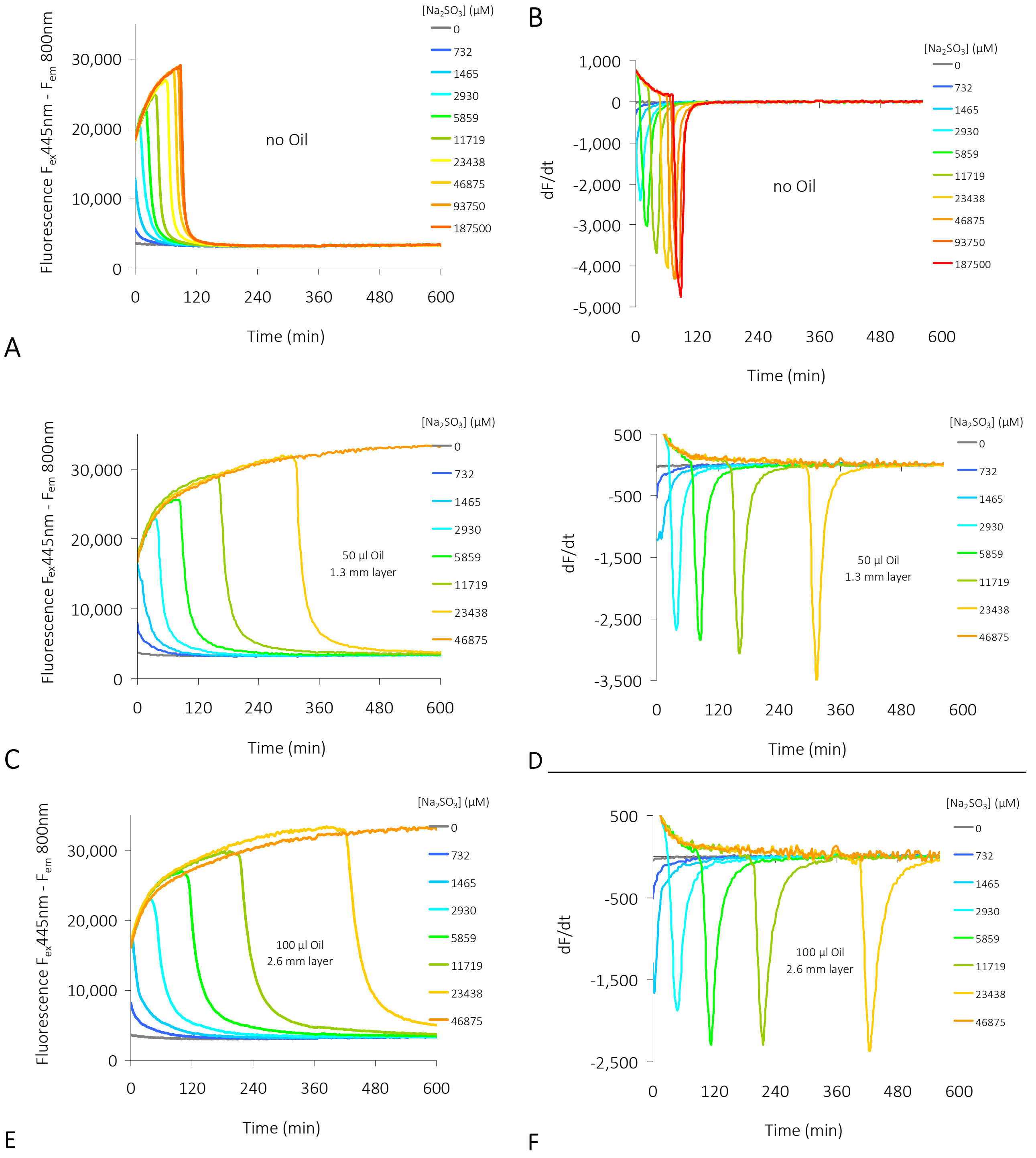


**Figure SI 6 $O_2$ ingress reduced with oil.** MTP wells with immobilized Pd-Benzoporphyrin (**2**) were filled with 200 µl solutions of diverse $[Na_2SO_3]$ as indicated in the legends and fluorescence was recorded over several hours to monitor $O_2$ ingress. **A, B**: wells uncovered; **C, D**: wells layered with 50 µl (≈ 1.3 mm) mineral oil. Reduction of $O_2$ ingress by a factor of 3.9. **E, F**: wells layered with 100 µl (≈ 2.6 mm) mineral oil. Reduction of $O_2$ ingress by a factor of 5.6. Assay conditions: 35°C, wells filled with 200 µl, $[NaSO_3]$ as indicated in the inset legends, sample rate ≈ 1/3 $min^{-1}$. The time derivatives in **B, D, F** are calculated from the data presented in **A, C, E**, respectively.

The diffusion process is clearly different when the probed aqueous phase is overlaid with oil because there is no direct contact between the atmosphere and the probed aqueous phase. This is seen when the quantity of $O_2$ consumed is plotted over the time needed to convert the provided amount of sulfite to sulfate (**Fig. SI 7**). As seen from **Fig. 7** in the main paper, oxygen consumption non-linearily correlates with time and increases exponentially when atmospheric $O_2$ has direct access to the probed solution. With an oil barrier, however, this process is slowed down and appears linear (**Fig. SI 7**).

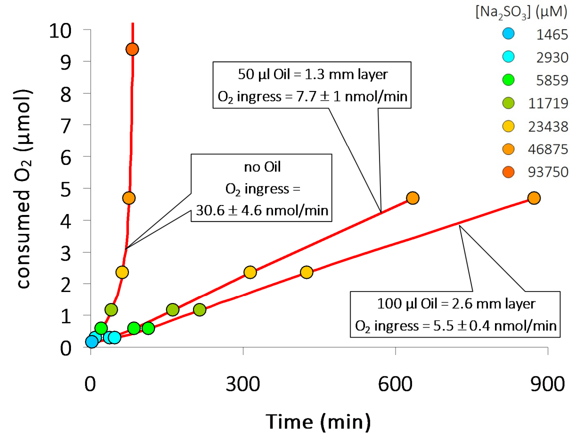


**Figure SI 7** The effect of oil on the $O_2$ ingress. The quantity of consumed $O_2$ as calculated from the given amount of sulfite is plotted over the time needed to convert the applied sulfite to sulfate. This allows the calculation of the $O_2$ ingress with and without oil layered onto 200 µl of $Na_2SO_3$ solution (concentrations in inset legend). A reduction of $O_2$ ingress by factors of 3.9 and 5.6 are achieved with 50 µl (≈ 1.3 mm) and 100 µl (≈ 1.3 mm) of mineral oil, respectively.

### $O_2$ ingress from residual air volume entrapped under the sealing film

Self adhesive $O_2$ impermeant films are effective to prevent undesired $O_2$ ingress into the solution to be probed for [$O_2$] in MTPs. However, when sealing the MTP, air is entrapped and the $O_2$ in the residual volume may disturb the experiment. Therefore, we measured $O_2$ ingress as described above (**Fig. 7** in the main text) in MTPs loaded with diverse volumes of liquid and sealed with polyester film. The sealed wells contained various amounts of $Na_2SO_3$ to determine speed and extent of the effect.

The summarized data (**Fig. SI 8**) show that the consumption time increases by a factor of 10 to 100 when the residual air volume in the wells is reduced from 350 µl to 100 µl. In conclusion, the undesirable effect can be drastically reduced, when the volume of entrapped air below the sealing film is kept as small as possible in each well. Some time-course data used to calculate the consumption times in **Fig. SI 8** are depicted in **Fig. SI 9**.

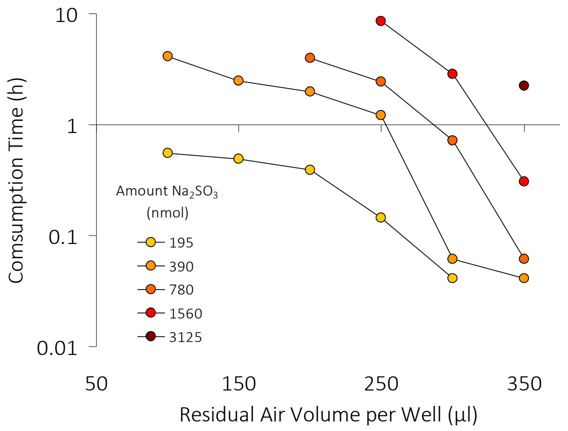


**Figure SI 8** The dependence of $O_2$ consumption on residual entrapped air volumes. Consumption times needed to oxidise various quantities of sulfite in dependence on the volume of air entrapped by the sealing film are depicted. The data were calculated from the experiment presented in **Fig. SI 9** and plotted over the entrapped air volume. The amount of $Na_2SO_3$ is given by the the inset legend.

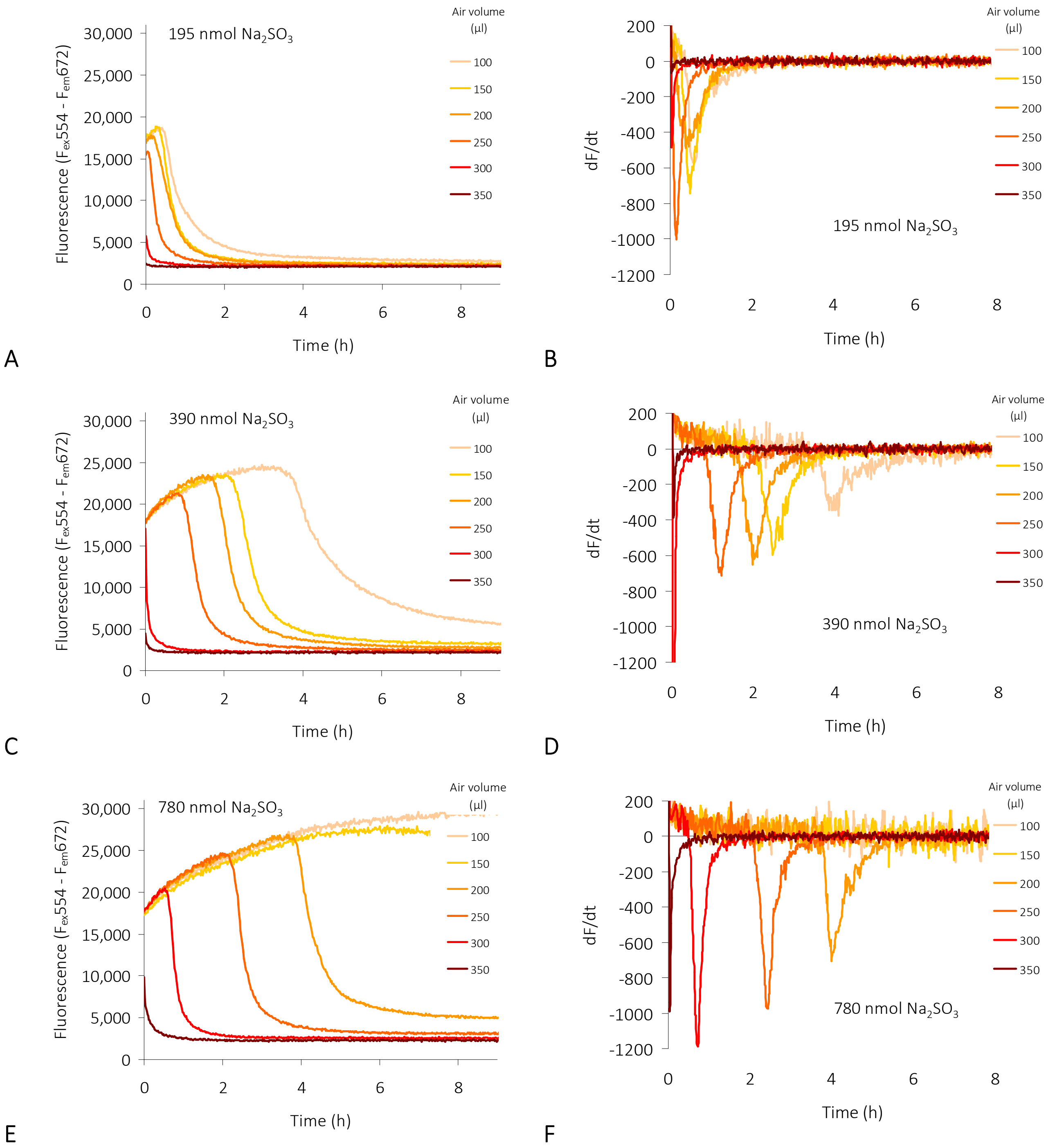


**Figure SI 9** **$O_2$ ingress observed with different entrapped air volumes in sealed wells containing various amounts of sulfite.** Wells were filled with the amount of sulfite indicated dissolved in $H_2O$ and sealed. The total well volume is 400 µl. The liquid volumes result from 400 µl minus the given residual air volume (inset legends). **A, C, E**: Pd-Fluoroporphyrin fluorescence; **B, D, F**: time derivatives calculated from data shown in opposite panel (A, C, E). Experimental conditions: 27°C; Sample rate = 4/5·$min^{-1}$.

### $O_2$ diffusion into and through plastic material.

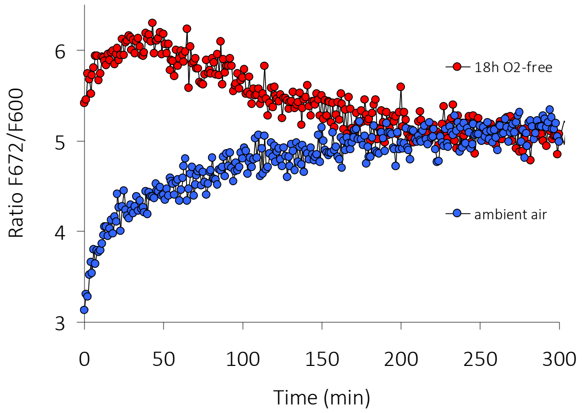


**Figure SI 10 $O_2$ absorption by and diffusion through the MTP plastic material.** A MTP was incubated overnight together with $O_2$-absorber in an $O_2$-free atmosphere to remove all $O_2$ bound to the plastic material. Wells were filled with sulfite solution and the fluorescence ratio F672/F600 was monitored for several hours (red dots). The same experiment with a MTP not pre-incubated and stored at ambient air was performed as a control (blue dots). Conditions: 25°C; sample rate =1 $\cdot min^{-1}$; 250 µl $[Na_2SO_3]$ = 300 mM.

Wells of MTPs incubated overnight in an $O_2$ free atmosphere in presence of $O_2$-absorber were filled with $Na_2SO_3$ solution and fluorescence ratios were recorded for several hours. Two different effects can be seen (**Fig. SI 10** red dots). First, the $O_2$ is consumed by the sulfite oxidation reaction (**Eq. 5** in the main text) and fluorescence ratio increases within the first 40 min due to the $[O_2]$ decrease. Second, this process of $O_2$ consumption is superseded by the diffusion of $O_2$ from the ambient atmosphere into and through the $O_2$-free MTP plastic material, leading to a ratio decrease and finally to an equilibrium at around R = 5, where $O_2$ ingress equals $O_2$ consumption by sulfite. The control data (**Fig. SI 10** blue dots) show, that the plastic is already $O_2$ saturated and the same equilibrium level is finally reached.

### pH-dependence of NMPs embedded in PS

PS is naturally not proton conductive. It lacks the chemical groups and residues necessary for facile proton transport. However, sulfonated polystyrene (S-PS), where sulfonic acid groups are attached to the molecular polystyrene backbone, exhibits proton conductivity [7–9]. This is because the sulfonic acid groups can dissociate, releasing protons ($H^+$) which are mobile and thus pass on through the material.

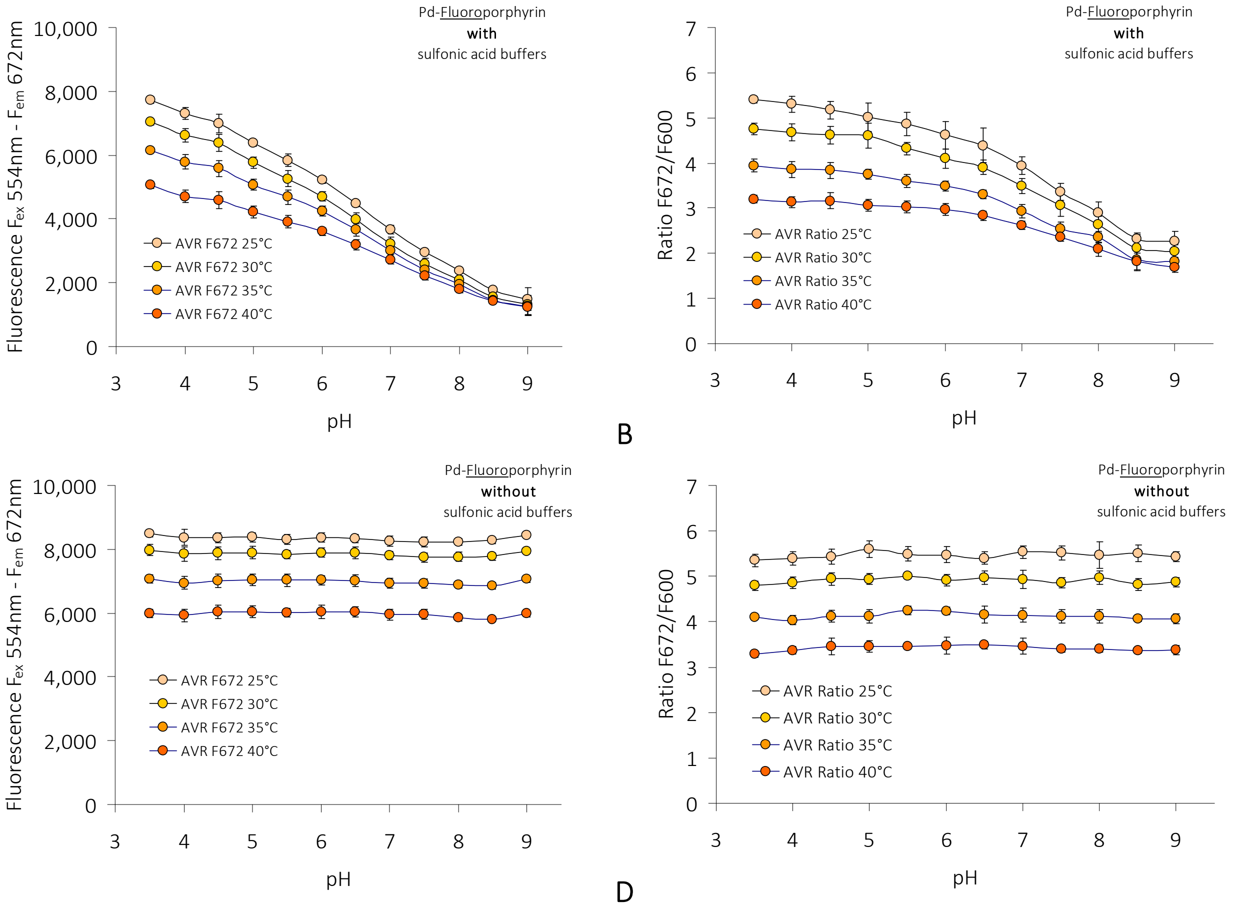


**Figure SI 11 pH-dependence of Pd-Fluoroporphyrin fluorescence emission under anoxia.** MTP-wells coated with **1** were filled with 200 µl of a mix of organic or mineral buffer and topped with 50 µl of $Na_2SO_3$ stock (1M), sealed with sealing film and incubated in $O_2$-free atmosphere for 2 days at RT. A: Fluorescence emission from **1** at various pH (x-axis) and under diverse temperatures (inset legend). pH was adjusted in a buffer mix containing various organic sulfonic acids. B: Fluorescence ratios calculated from data in A and F600 recorded in parallel. C and D: Same as in A and B, however pH was adjusted in a mix of mineral buffers. Assay conditions for A and B - mix of organic buffers containing sulfonic acids: Citric acid, MES, HEPES, TAPS; 50 mM each; adjusted with KOH to desired pH. Assay conditions for C and D: Mix of citric acid and inorganic buffers without sulfonic acids: Citric acid, $KH_2PO_4$, $K_2HPO_4$, $K_2CO_3$, 50 mM each; adjusted with KOH to desired pH. AVR of n = 7; error bars represent SD.

To check this effect, MTP wells doped with **1** or **2** were filled with a buffer mix adjusted to a series of different pH values ($3.5 \leq pH \leq 9$) and fluorescence was measured after 2 days incubation in $O_2$-free atmosphere. Sulfonic acid buffers cause a pronounced pH dependence (Figs. SI 11 A, B; 12 A, B), revealing a $H^+$-permeability of PS, whereas mineral buffers do not show this effect (Fig.SI 11 5 C, D; 12 C, D). Fluorescence emission of **1** and **2** embedded in PS becomes pH-dependent when the plastic is in contact with sulfonic acids. This effect is reversible since it disappeared when MTPs were thoroughly washed and rinsed with distilled water and refilled with mineral buffer (data not shown).

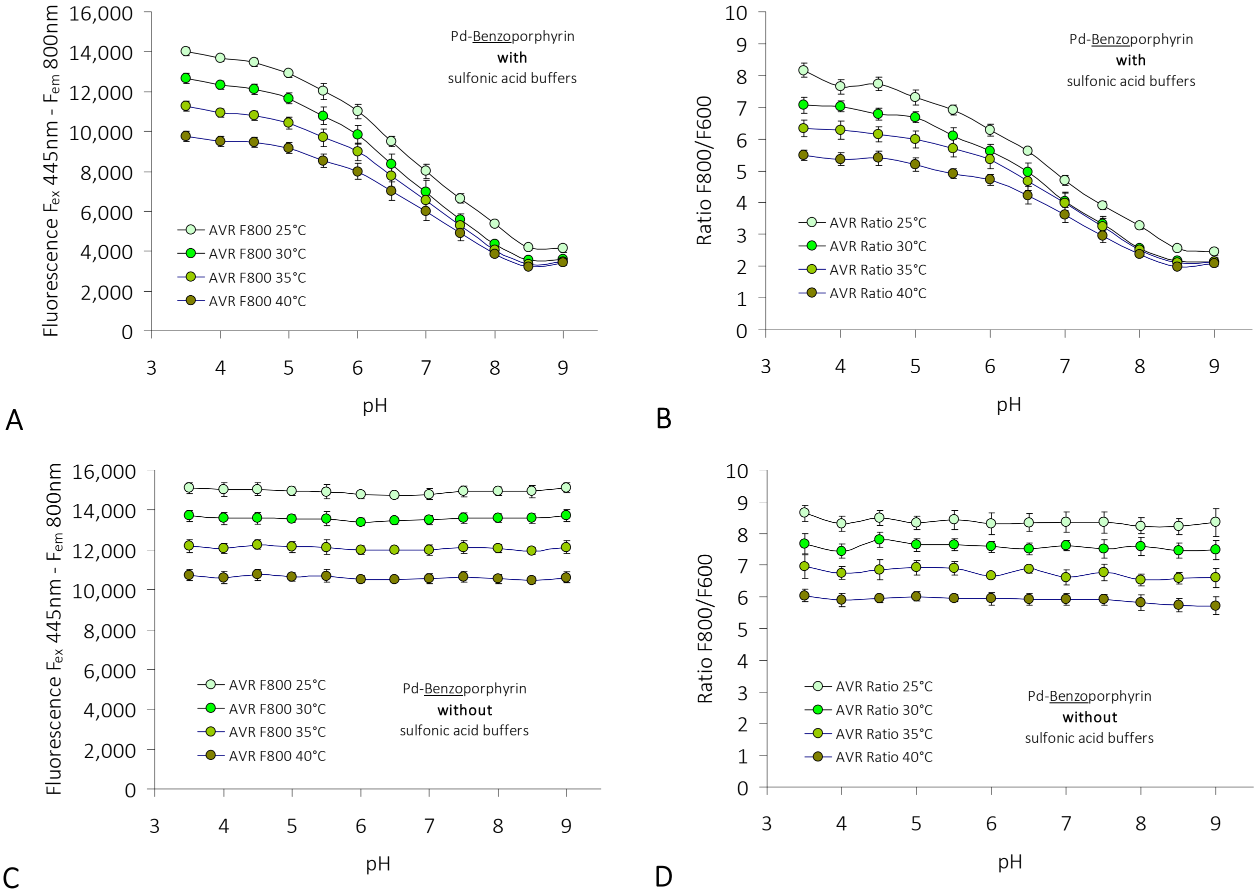


Figure SI 12 pH-depencence of Pd-Benzophorphyrin fluorescence emission under anoxia. MTP-wells coated with **2** were filled with 200 µl of a mix of organic or mineral buffer and topped with 50 µl of $Na_2SO_3$-stock (1M), sealed with sealing film and incubated in $O_2$-free atmosphere for 2 days at RT. A: Fluorescence emission from **2** at various pH (x-axis) and under diverse temperatures (inset legend). pH was adjusted in a buffer mix containing various organic sulfonic acids. B: Fluorescence ratios calculated from data in A and F600 recorded in parallel. C and D: Same as in A and B, however pH was adjusted in a mix of mineral buffers. Assay conditions for A and B - mix of organic buffers containing sulfonic acids: Citric acid, MES, HEPES, TAPS; 50 mM each; adjusted with KOH to desired pH. Assay conditions for C and D: Mix of citric acid and inorganic buffers without sulfonic acids: Citric acid, $KH_2PO_4$, $K_2HPO_4$, $K_2CO_3$, 50 mM each; adjusted with KOH to desired pH. AVR of n = 7; error bars represent SD.

Table SI 1 Chemicals

| | |
|---|---|
| Brain-Heart-Infusion-Broth | Roth #X916; |
| Catalase (CAS 9001-05-2) 5 MU 40.000 U/mg Protein | Roth #6025; |
| Chloroform | Roth # T901 or Sigma-Aldrich-Merck #288306; |
| Citric acid (CAS 77-92-9) | Sigma-Aldrich-Merck #27487; |
| di-Potassium hydrogen phosphate (CAS No. 7758-11-4) | Roth # T875; |
| di-Sodium hydrogen phosphate (CAS 7558-79-4) | Roth # T876; |
| Hydrogen peroxide (CAS 7722-84-1) | Roth #8070; |
| Hydroxyethylpiperazine-ethane sulphonic acid (HEPES Buffer; CAS 7365-45-9) | Roth #9105; |
| Kanamycin sulfate (CAS 25389-94-0) | Roth #T832; |
| Laccase from *Trametes versicolor* (CAS 80498-15-3) | Sigma-Aldrich-Merck #51639; |
| LB-Medium (Luria/Miller) | Roth # X969; |
| Magnesium Chlorid Hexahydrate (CAS 7791-18-6) | Sigma-Aldrich-Merck #1.05832; |
| Mineral Oil | Roth #HP50; |
| Morpholino-ethane sulphonic acid (MES Buffer; CAS 4432-31-9) | Roth #4256; |
| Murashige & Skoog Medium | Duchefa # M0222; |
| Oxygen Absorber | Midukit #X001RVBI4L; |
| Palladium (II) 5,10,15,20-(tetrapentafluorophenyl) - porphyrin (CAS 72076-09-6) | PorphyChem #03211525; |
| Palladium (II) 5,10,15,20-(tetraphenyl) - tetrabenzoporphyrin (CAS 119654-64-7) | PorphyChem #03323825; |
| Petroleum-Benzine 40-60 | Roth #9320; |
| Potassium carbonate (CAS 584-08-7) | Roth #7956; |
| Potassium chloride (CAS 7447-40-7) | Roth #6781; |
| Potassium dihydrogen phosphate (CAS 7778-77-0) | Roth #3904; |
| Potassium Hydroxide (CAS 1310-58-3) | Roth #6751; |
| Pyrogallol (CAS 87-66-1) | Roth #4364; |
| Sodium hydroxide (CAS 1310-73-2) | Roth #6771; |
| Sodiumchloride (CAS 7647-14-5) | Roth #3957; |
| Tris(hydroxymethyl)-amino methane (TRIS Buffer; CAS No. 77-86-1) | Roth # AE15; |
| Tris(hydroxymethyl)-amino butane sulfonic acid (TABS Buffer; CAS 54960-65-5) | Sigma-Aldrich-Merck #T1302; |
| Tris(hydroxymethyl)-amino-propane sulphonic acid (TAPS Buffer; CAS No. 29915-38-6) | Roth #6982; |
